# Calcium-induced calcium release in noradrenergic neurons of the locus coeruleus

**DOI:** 10.1101/853283

**Authors:** Hiroyuki Kawano, Sara B. Mitchell, Jin-Young Koh, Kirsty M. Goodman, N. Charles Harata

## Abstract

The locus coeruleus (LC) is a nucleus within the brainstem that consists of norepinephrine-releasing neurons. It is involved in broad processes including autonomic regulation, and cognitive and emotional functions such as arousal, attention and anxiety. Understanding the mechanisms that control the excitability of LC neurons is important because they innervate widespread regions of the central nervous system. One of the key regulators is the cytosolic calcium concentration ([Ca^2+^]_c_), the increases in which can be amplified by calcium-induced calcium release (CICR) from the intracellular calcium stores. Although the electrical activities of LC neurons are regulated by changes in [Ca^2+^]_c_, the extent of CICR involvement in this regulation has remained unclear. Here we show that CICR hyperpolarizes acutely dissociated LC neurons of the rat brain and demonstrate the pathway whereby it does this. When CICR was activated by extracellular application of 10 mM caffeine, LC neurons were hyperpolarized in the current-clamp mode of the patch-clamp recording, and the majority of neurons showed an outward current in the voltage-clamp mode. This outward current was accompanied by an increase in membrane conductance, and its reversal potential was close to the K^+^ equilibrium potential, indicating that it is mediated by the opening of K^+^ channels. The outward current was generated in the absence of extracellular calcium and was blocked when the calcium stores were inhibited by applying ryanodine. Pharmacological experiments indicated that the outward current was mediated by Ca^2+^-activated K^+^ channels of the non-small conductance type. Finally, the application of caffeine led to an increase in the [Ca^2+^]_c_ in these neurons, as visualized by fluorescence microscopy. These findings delineate a mechanism whereby CICR suppresses the electrical activity of LC neurons, and indicate that it could play a dynamic role in modulating the LC-mediated noradrenergic tone in the brain.

## INTRODUCTION

The locus coeruleus (LC) is a nucleus within the dorsal pontine region of brainstem, and serves as the major source of noradrenergic input to various regions throughout the central nervous system (Szabadi, 2013; Schwarz et al., 2015; Uematsu et al., 2017; Chandler et al., 2019; Zerbi et al., 2019). It has been implicated in the regulation of diverse functions of both the central nervous system and the peripheral autonomic nervous systems, including the activity of the sympathetic nervous system, arousal, attention, memory, sensory information processing, anxiety, sleep and pain sensation (Sara and Bouret, 2012; Sara, 2015; Aston-Jones and Waterhouse, 2016; Chandler, 2016; Takeuchi et al., 2016; Manella et al., 2017; Totah et al., 2018; Rodenkirch et al., 2019; Zerbi et al., 2019). Consistent with these roles, abnormal regulation of LC neuron activity has been implicated in a wide variety of neurological and psychiatric disorders, including autonomic dysfunction (Vermeiren and De Deyn, 2017), Parkinson’s disease (Espay et al., 2014; Schapira et al., 2017; Vermeiren and De Deyn, 2017; Peterson and Li, 2018), Alzheimer’s disease (Giorgi et al., 2017; Peterson and Li, 2018), dystonia (Hornykiewicz et al., 1986; McKeon et al., 1986), major depression (Fan et al., 2018), anxiety disorders (McCall et al., 2017), bipolar disorder (Cao et al., 2018), and migraine (Vila-Pueyo et al., 2019). Thus LC activity is crucial to the functions of the nervous system, and a deeper understanding of its regulation is expected to have an impact on the research of a variety of disorders.

The cytosolic calcium concentration ([Ca^2+^]_c_) is an important determinant of neuronal activities (Sanchez-Padilla et al., 2014; Matschke et al., 2015; Rawal et al., 2019) and is modified by the mechanism, Ca^2+^-induced Ca^2+^ release (CICR) from intracellular Ca^2+^ stores. Triggered by an initial small increase in [Ca^2+^]_c_, for instance the Ca^2+^ influx from the extracellular space into the cytosol, this mechanism further increases [Ca^2+^]_c_ by enabling Ca^2+^ release from the stores through Ca^2+^-permeable ryanodine receptors (Verkhratsky, 2005; Endo, 2009; Meissner, 2017; Popugaeva et al., 2017). The Ca^2+^ signal amplification by CICR has been demonstrated in numerous types of neurons of both central and peripheral nervous systems, and plays a role in propagating the [Ca^2+^]_c_ rises that control neuronal membrane excitability, synaptic plasticity and gene expression (Verkhratsky, 2005; Bardo et al., 2006). One consequence of CICR is a change in Ca^2+^-sensitive ionic conductance across the plasma membrane. Caffeine and its analog theophylline are known to directly promote CICR (Kuba, 1980; McPherson et al., 1991; Friel and Tsien, 1992; Usachev et al., 1993; Llano et al., 1994; Kano et al., 1995; Meissner, 2017) by sensitizing ryanodine receptors and thus lowering the threshold of CICR (Albrecht et al., 2002; Kong et al., 2008). Both compounds increase [Ca^2+^]_c_ (Smith et al., 1983; Neering and McBurney, 1984) and induce a Ca^2+^-activated K^+^ current (Kuba, 1980; Munakata and Akaike, 1993). Induction of the Ca^2+^-activated K^+^ currents by CICR is also widespread (Verkhratsky, 2005), occurring in, e.g. sympathetic neurons of the bull-frog (Akaike and Sadoshima, 1989; Marrion and Adams, 1992), neurons of dorsal motor nucleus of the vagus of guinea-pig (Sah and McLachlan, 1991), and hippocampal CA1 pyramidal neurons of the rat (Uneyama et al., 1993; Garaschuk et al., 1997). Physiological induction of Ca^2+^-activated K^+^ currents, e.g. by membrane depolarization followed by Ca^2+^ influx, leads to a reduction in the frequency of action potential firing (Stocker, 2004; Salkoff et al., 2006; Matschke et al., 2018) and in network excitability (Li et al., 2019).

Despite the importance of neuronal CICR, the detailed properties of CICR in LC neurons have been largely unexplored, except for the following two lines of research. First, the membrane hyperpolarization following action potentials (after-hyperpolarization) has been shown to depend on increases in [Ca^2+^]_c_ (Aghajanian et al., 1983). It is composed of two phases. The fast phase is activated by the Ca^2+^ influx through voltage-dependent Ca^2+^ channels and mediated by opening of Ca^2+^-activated K^+^ channels of the small-conductance type (SK) without an involvement of CICR (Aghajanian et al., 1983; Osmanovic et al., 1990; Matschke et al., 2018). The slow phase was suppressed by reagents that block Ca^2+^ release from intracellular Ca^2+^ stores (thus demonstrating the presence of CICR in LC neurons) and was mediated by Ca^2+^-activated K^+^ channels, yet their properties have not been identified. Second line of research revealed that when CICR is stimulated by the application of caffeine, an outward current is generated in LC neurons (Murai et al., 1997; Ishibashi et al., 2009).

This finding suggested that K^+^ channels may be opened, as is the case during the slow phase of after-hyperpolarization. These studies showed that the CICR takes place in LC neurons. However, neither the [Ca^2+^]_c_ increase by CICR, nor the dynamics of this increase, has been visualized in LC neurons, and thus the direct proof that the CICR contributes to the increase in [Ca^2+^]_c_ has been lacking. It also remains unclear how direct activation of CICR influences the electrical properties of LC neurons, for example, the effects on membrane potential, the type and kinetics of activated ion channels.

Here we addressed these questions in rat LC neurons. In order to observe CICR in isolation from other [Ca^2+^]_c_-elevating mechanisms such as synaptic transmission and glial effects on neurons, we acutely dissociated the LC cells. After morphologically confirming that those cells are noradrenergic neurons, we demonstrate that the major effect of caffeine was to hyperpolarize the membrane potential, thereby suppressing the generation of action potentials. We also show that the application of caffeine released Ca^2+^ from the intracellular Ca^2+^ stores, increased [Ca^2+^]_c_ in dendrites and soma, and opened the Ca^2+^-activated K^+^ channels of non-SK type. The degree of Ca^2+^ store filling seems to be associated with the activation kinetics of the K^+^ current.

## MATERIALS AND METHODS

### Neuron preparation

Animal care and use procedures were approved by the University of Iowa’s Institutional Animal Care and Use Committee, and performed in accordance with the standards set by the PHS Policy and *The Guide for the Care and Use of Laboratory Animals* (NRC Publications) revised 2011. Neurons were acutely dissociated from the rat LC according to the following protocol. Nine- to fourteen-day-old Sprague-Dawley rats (Charles River Laboratories International, Inc., Wilmington, MA, USA) were decapitated under intraperitoneal anesthesia with ketamine (60-100 mg/kg) and xylazine (10-15 mg/kg). The brain was quickly removed from the skull, transferred to ice-cold incubation medium II (see below for solution compositions) saturated with 95% O_2_ - 5% CO_2_ gas, and sliced at a thickness of 400 µm (VT1200 S microslicer, Leica Biosystems, Buffalo Grove, Illinois, USA). Following 30-60 min of incubation in incubation medium I at room temperature, the slices were treated with pronase (1 mg / 4-6 ml) (Calbiochem-MilliporeSigma, Burlington, MA, USA) in incubation medium I at 31°C for 30-50 min. Thereafter they were left in enzyme-free medium I at room temperature for 30 min. Similar results were obtained when the enzyme treatment procedure was replaced by dispase (10,000 Unit / 6-8 ml) (Calbiochem-MilliporeSigma) at 31°C for 40-60 min, followed by leaving the slices in enzyme-free incubation medium I for 1.5-2.5 hours at room temperature. After these treatments were completed, the slices were transferred to the standard external solution in a 35-mm culture dish (BD Falcon 353001, BD, Franklin Lakes, NJ, USA) coated with silicone. The LC region was identified under a binocular stereomicroscope (SMZ645, Nikon, Melville, NY, USA), and was removed by micro-punching with a blunt syringe needle. For electrophysiology experiments, the micro-punched specimens were transferred to a fresh culture dish (BD Falcon 353001) containing the standard external solution, and mechanically triturated using a fire-polished Pasteur pipette. The dissociation procedure was monitored by phase-contrast microscopy (Eclipse TS100, Nikon). For imaging experiments, the micro-punched specimens were triturated by the same procedure, but in a different type of culture dish (0.17-mm-thick glass coverslip bottom, Delta T culture dish, Bioptechs, Butler, Pennsylvania, USA). The dissociated neurons adhered to the bottom of the dish within 30 min.

### Phase-contrast microscopy

Acutely dissociated cells were observed under a microscope (Eclipse Ti-E, Nikon), using a 20x (CFI S Plan Fluor ELWD; numerical aperture 0.45) or 40x air objective lens (CFI S Plan Fluor ELWD; numerical aperture 0.60) with Köhler illumination for transmitted light microscopy (Centoze and Pawley, 2006). In all experiments in this study, we used live cells under good conditions or fixed cells that would have been under good condition at the time of fixation, based on the following morphological characteristics. Soma is phase-bright and the soma and dendrite show a smooth cell surface. The dendrites are often phase-bright, but occasionally phase-dark. Morphological characteristics of dead or unhealthy cells are: a phase-dark soma and dendrites, with the visible nucleus, and a non-smooth soma surface and occasional beading of dendrites. These features were confirmed by live-dead staining (Supplementary Figure S1). The area of soma was measured using ImageJ (Rasband, W.S., NIH, Bethesda, MD, USA).

**Figure 1.**
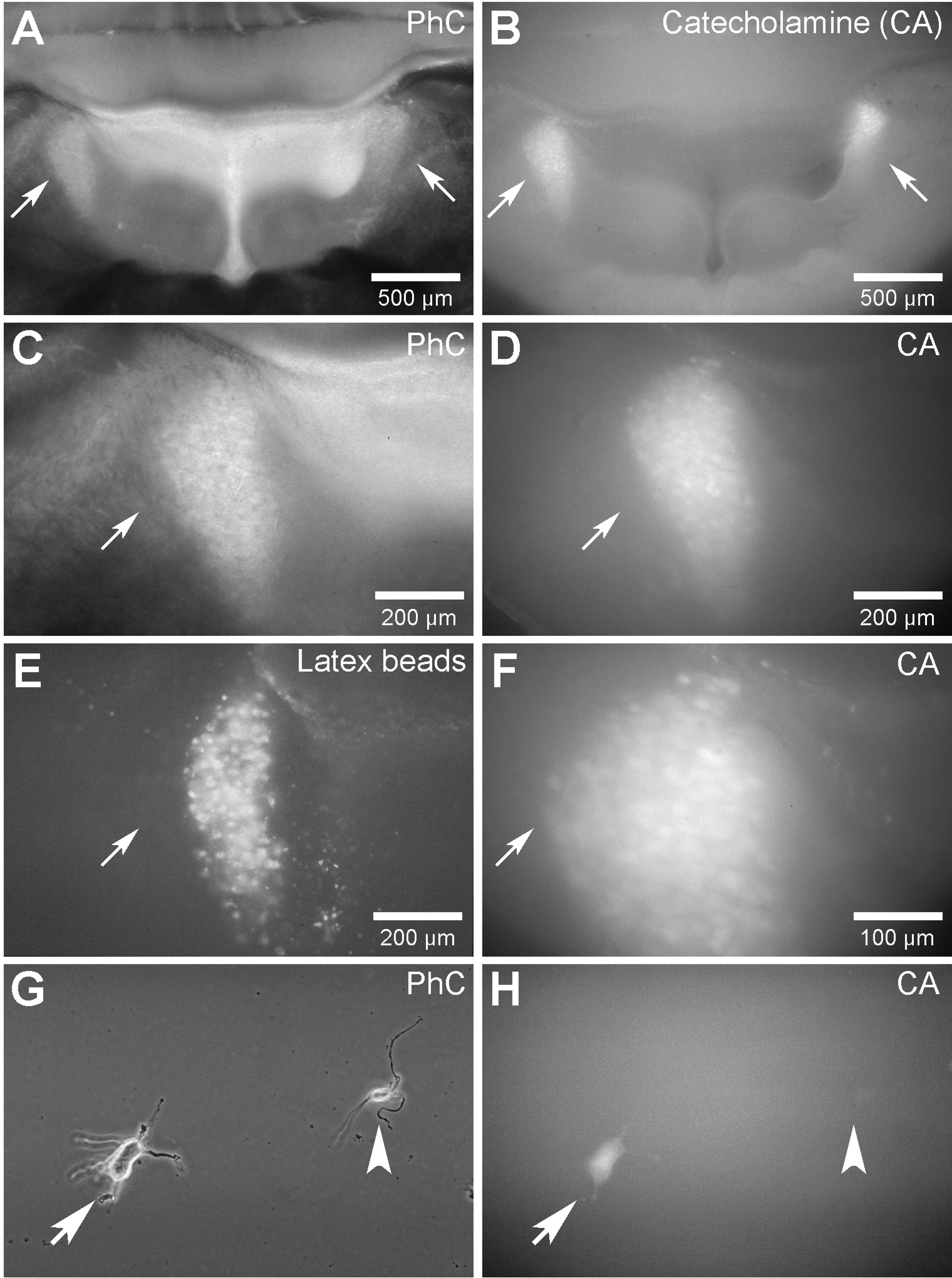
Identification of the locus coeruleus (LC) in brain slices and acutely dissociated LC neurons. **(A,B)** Single slice of the brainstem area at the level of the fourth ventricle, viewed by **(A)** phase-contrast (PhC) imaging and **(B)** fluorescence imaging of catecholamines (CA). Arrows in **(A)** and **(B)** point to bilateral regions of the LC. Although the shapes of these bilateral regions appear asymmetrical in this figure, this is likely due to slicing at an oblique angle. **(C, D)** Higher magnification views of images shown in **(A)** and **(B)**, respectively, focusing on left side of section. **(E)** Fluorescence image of a brain slice at a similar level as in **(A)**, five days after bilateral injection of fluorescent latex beads into the cerebral cortex. **(F)** Higher magnification view of image shown in **(D)**. **(G, H)** Cells acutely dissociated from the LC region, viewed by phase-contrast imaging **(G)** and fluorescence imaging of catecholamines **(H)**. Note that the large neuron was positive for catecholamine (arrow, positive staining visible in soma), whereas the small neuron was negative (arrowhead).

### Electrical recordings

Electrical recordings were carried out by the nystatin-perforated mode of the whole-cell patch-clamp recording. This mode prevents diffusion-based exchange of second messenger-related molecules between the patch pipette and the cell interior, thus maintaining the intracellular environment (Horn and Marty, 1988; Akaike and Harata, 1994; Harata et al., 1997). Patch-pipettes were fabricated from borosilicate glass tubes (1.5-mm outer diameter; G-1.5, Narishige International USA, East Meadow, NY, USA), in two stages and using a vertical pipette puller (PB-7, Narishige International USA). The resistance of the recording electrode was 5-7 MΩ. Liquid junction potentials of 3-4 mV were used to calibrate the holding potential. The current and voltage were measured with a patch-clamp amplifier (Axopatch 200B, Molecular Devices, Sunnyvale, CA, USA), which was controlled by either the pCLAMP software (Molecular Devices), or a custom-written software based on the graphical programming environment LabVIEW (National Instruments, Austin, TX, USA). The membrane currents were filtered at 7 kHz and were digitized at 25 kHz. The electrical system was mounted on an inverted microscope (Eclipse Ti-E, Nikon).

Ramp voltage command consisted of a linear hyperpolarizing and then depolarizing voltage command of 51 mV with a frequency of 0.4 Hz (40.8 mV/sec). The voltage-dependent Na^+^ and Ca^2+^ currents were suppressed during the ramp-wave studies, by adding 1 µM tetrodotoxin (Tocris Bioscience, Ellisville, MO, USA) and 10 µM LaCl_3_ to the external solutions. Except for ramp voltage commands, the holding potential was set at −44 mV for the voltage-clamp recording (Munakata and Akaike, 1993).

### Identification of LC based on retrograde tracing

The LC region was visualized by injecting rat brains with fluorescent microspheres. These reagents are taken up by axons and transported to neuronal soma, and thus serve as retrograde tracers of axons. Specifically, red-fluorescent FluoSpheres (F8793), Molecular Probes-Thermo Fisher Scientific, Walthamm, MA, USA) (Maruyama and Ohmori, 2006) were injected into the cerebral cortices of the animals (n=3), through 8 holes made bilaterally in the skull under anesthesia. Three to five days after injection, the animals were euthanized and the brains were isolated as described above. 400 µm-thick brain slices were cut and examined by widefield fluorescence microscopy (IX70, Olympus, Center Valley, PA, USA) (“G” filter set; excitation, 480-550 nm; dichroic mirror, 570 nm; emission, 590-nm long pass). A similar retrograde labeling of LC was reported with a fluorescent dye (DiI) injected into spinal cord (Zhang et al., 2007).

### Identification of LC based on catecholamine fluorescence

Classical formaldehyde-glutaraldehyde-induced catecholamine fluorescence was also used to identify the LC region and the neurons of this nucleus within the brain (Furness et al., 1977a; Furness et al., 1977b; Jiang et al., 1994). To this end, rats were anesthetized, transcardially perfused with phosphate-buffered saline (PBS; 70011044; GIBCO-Thermo Fisher Scientific, Waltham, MA, USA) and perfusion-fixed with 4% paraformaldehyde and 0.5% glutaraldehyde (Electron Microscopy Sciences, Hatfield, PA, USA) in PBS at room temperature. The brains were dissected out and post-fixed in the same solution at room temperature for 3-6 hours. The brain slices containing the LC region were then sectioned in PBS, at a thickness of 400 µm. They were immediately examined using an inverted microscope fitted with an widefield fluorescence system (IX70, Olympus) (“V” filter set; excitation, 380-430 nm; dichroic mirror, 455 nm; emission, 455-nm long pass).

Acutely dissociated neurons were tested for noradrenergic properties according to the following procedure. Live, dissociated neurons that had settled on coverslips (Delta T culture dish, Bioptechs) were rinsed twice with PBS. These neurons were then fixed with 4% paraformaldehyde and 0.5% glutaraldehyde in PBS for 3-6 hours at room temperature, rinsed 2-3 times with PBS, and imaged by widefield fluorescence microscopy. To assess the specificity of the formaldehyde-glutaraldehyde-induced catecholamine fluorescence, we treated the dissociated and stained neurons with 0.1% sodium borohydride in PBS at room temperature for 20 min, a treatment known to abolish catecholamine fluorescence while leaving the non-specific fluorescence intact (e.g. autofluorescence) (Corrodi et al., 1964). Indeed, catecholamine fluorescence was significantly reduced (data not shown).

### Characterization of LC neurons by immunocytochemistry

Immunocytochemistry was performed to confirm and extend the catecholamine fluorescence data, by staining the neuronal dendrite marker, microtubule-associated protein 2 (MAP2) (Craig and Banker, 1994) and the noradrenergic marker, dopamine β-hydroxylase (DBH), the enzyme that synthesizes norepinephrine (Foote et al., 1983; Sanchez-Padilla et al., 2014; Schmidt et al., 2019). Dissociated neurons were fixed with 4% paraformaldehyde and 4% sucrose in the standard external solution at 4°C for 30 min. After rinses with the standard external solution twice for 5 min each, the neurons were permeabilized and blocked with 0.4% saponin and 2% normal goat serum (Jackson ImmunoResearch Laboratories, West Grove, PA, USA) in PBS at room temperature for 60 min. They were incubated with a polyclonal rabbit anti-MAP2 antibody (AB5622, Chemicon-MilliporeSigma, Burlington, MA, USA) and a monoclonal mouse anti-DBH antibody (MAB308, Chemicon-MilliporeSigma) (both diluted 400-fold) in permeabilization-block solution, overnight at 4°C. Following rinses in PBS 3 times for 7 min each, the neurons were incubated in goat anti-rabbit secondary antibody conjugated with quantum dot 525 (Molecular Probes-Thermo Fisher Scientific; diluted 50-fold) and goat anti-mouse secondary antibody conjugated with quantum dot 605 (Molecular Probes-Thermo Fisher Scientific; diluted 50-fold) in permeabilization-block solution, at room temperature for 60 min. They were rinsed in PBS at least 5 times for 20 min each. The neurons were then observed under an inverted microscope (TE300, Nikon) with a CCD camera (DP71, Olympus). The excitation light was obtained from a light source (X-Cite 120, Lumen Dynamics Group Inc., Ontario, Canada). Signals from quantum dot 525 were obtained with excitation and emission filters of 470/40, 525/50 nm, and those from quantum dot 605 were obtained with 545/30, 610/75 nm. The neurons were analyzed for their size and the fluorescence intensity of staining, using ImageJ. The fluorescence intensity of a cell was measured by defining a region of interest (ROI) around the perimeter of the soma in a phase-contrast image, and averaging the pixel intensity within the ROI in a fluorescence image. The intensity was corrected by subtracting the fluorescence intensity outside the cells (acellular background). The images were subjected to 3-dimensional deconvolution.

### Confocal Microscopy

Confocal laser scanning microscopy was carried out using a Noran Odyssey system (Noran Instruments, Middelton, Wisconsin, USA) adapted to an inverted microscope (Diaphot TMD 200, Nikon). The 60x objective lens (Plan Apo, water immersion, NA = 1.2, Nikon) was used. The fluorophores were excited using a 488-nm line ion-argon laser, and the emission longer than 515 nm was collected. The system used an acousto-optical deflector to provide adjustable scanning rates along the x-axis. The laser intensity was set at 30-40%. The fixed cells were scanned with a dwell time of 400.00 nsec, and 16 frames were averaged to give a single image (0.47 images/sec). Pixel size was 0.12 x 0.12 μm (x-y dimension), and z spacing was 0.50 μm. The slit width for detection was set at 15 µm. All acquired images were saved in an 8-bit gray-scale format. The fluorescence intensity of a subcellular region was measured by placing a ROI in a maximum-intensity projection image, and averaging the pixel intensities using ImageJ. The intensity was corrected by subtracting the fluorescence intensity of the acellular background.

For [Ca^2+^]_c_ imaging, membrane-permeable, Fluo-3-acetoxymethylester (Fluo-3-AM) (Molecular Probes-Thermo Fisher Scientific) was used (Beauvais et al., 2016), because changes in path length and dye concentration were minimal within each imaging session (8 sec) (Cornell-Bell et al., 1990). Dissociated neurons were incubated in the standard external solution with 1 μM Fluo-3-AM and 0.001% cremophor EL at room temperature for 30-40 min. After careful washing with dye-free standard external solution, the neurons were used for experiments within 30 min to 2 hours. Neurons were scanned at 30 frames/sec with a dwell time of 100.00 nsec, and 2 frames were averaged to give a single image (15 images/sec or 67 msec/image). Pixel size was 0.15 x 0.15 μm (x-y dimension). The slit width for detection was set at 25 µm, which provides an axial resolution of <1.3 µm with the objective lens used. The neurons that emitted brightly with 20% laser intensity or less under resting conditions were discarded, because they usually responded poorly to caffeine application: thus the high resting level of [Ca^2+^]_c_ indicates poor condition of cell health. The plane of scanning was adjusted to allow observation of a dendrite at its broadest diameter. The present study was confined to neurons whose dendrites emerged from the bottom of the soma, close to the floor of the dish, and whose dendrites could be traced in a single plane. Neurons whose dendrites emerged above the bottom were excluded because it was not possible to image their dendrites in a single plane. For numerical analyses, we measured the average pixel intensity within ROIs as the raw fluorescence intensity (F). The intensity was corrected for the pre-response intensity inside the cell (baseline, F_0_) and the intensity outside the cell (acellular background, F_b_), using the following formula: (F−F_0_)/(F_0_−F_b_) = ΔF/(F_0_−F_b_). Numerical data were analyzed using custom-written software based on LabVIEW.

### Solutions

The ionic composition (all in mM) of the Incubation Medium I for room temperature and 31°C was: NaCl 124, KCl 5, KH_2_PO_4_ 1.2, NaHCO_3_ 24, CaCl_2_ 2.4, MgSO_4_ 1.3, glucose 10 and sucrose ∼16. Prior to the addition of sucrose, the osmolarity of the solution measured approximately 293 mOsm. Sucrose was added to increase the osmolarity to 310 mOsm. The ionic composition of the Incubation Medium II for 4°C was: NaCl 124, KCl 5, KH_2_PO_4_ 1.2, NaHCO_3_ 34, CaCl_2_ 2.4, MgSO_4_ 1.3 and glucose 10. The NaHCO_3_ concentration was increased to attain a correct pH. The osmolarity of the solution measured approximately 309 mOsm without sucrose. The pH of both incubation media was adjusted to 7.45 with 95% O_2_ - 5% CO_2_ gas. To obtain a stable pH value, the media were bubbled for a minimum of 30 min (typically ∼60 min). The ionic composition of the standard external solution was: NaCl 150, KCl 5, CaCl_2_ 2, MgCl_2_ 1, glucose 10 and *N*-2-hydroxyethylpiperazine-*N’*-2-ethanesulfonic acid (HEPES) 10. The 30 mM K^+^ solution was prepared from the standard external solution by replacing 25 mM NaCl with the same concentration of KCl. The ionic composition of the internal (patch-pipette) solution for nystatin-perforated patch recording was: KCl 150 and HEPES 10. The pH of the standard external and internal solutions was adjusted to 7.4 and 7.2, respectively, with tris(hydroxymethyl)aminomethane (Tris-OH). Nystatin was dissolved in acidified methanol at 10 mg/ml. The stock solution was dissolved in the internal solution just before use, at a final concentration of 100-200 µg/ml.

### Drug treatments

Chemical reagents were purchased from Sigma-Aldrich (St. Louis, MO, USA) unless mentioned otherwise. Pharmacological reagents were applied to live neurons by the “Y-tube” method, a fast application system that allows the external solution to be exchanged within 30 msec, with an average travel rate of ∼100 μm/msec near the cell (Harata et al., 1996; Kira et al., 1998; Harata et al., 1999; Iwabuchi et al., 2013). This method ensures that all parts of a single neuron are exposed to caffeine within the duration of a single image frame (67 msec).

## RESULTS

### Identification of LC Region in Rat Brain Slices, and of LC Neurons Following Acute Dissociation

Both the LC in brain slices and the acutely dissociated LC neurons were identified based on the classical catecholamine fluorescence. This approach takes advantage of the fact that exposure to a formaldehyde-glutaraldehyde mixture induces catecholamines, including the neurotransmitter norepinephrine, to fluoresce (**Figure 1**). Examination of brain slices by phase-contrast microscopy revealed semi-triangular, phase-lucent areas on either side of the brainstem at the level of the fourth ventricle (arrows in **Figure 1A,C**). The same areas fluoresced brightly, identifying this structure as the catecholamine-containing LC (arrows in **Figure 1B,D**). A high-power view of the same section made it possible to visualize the individual somata of stained neurons (**Figure 1F**). Among neurons acutely dissociated from the phase-lucent regions of unfixed brain slices (**Figure 1G**), those that were large were positive for catecholamine fluorescence, whereas those that were small were negative (**Figure 1H**, arrow vs. arrowhead, respectively).

To confirm the identification of LC in brain slices, we localized the LC region based on its innervation of broad regions of the CNS, in particular the cerebral cortex (Schwarz et al., 2015). This experiment involved injecting fluorescent latex beads into the cerebral cortex and following their distribution in brainstem slices after leaving time for their retrograde transport to neuronal soma. Three to five days after injection, the region that was translucent by phase-contrast microscopy was also populated by neurons that fluoresced intensely due to the presence of latex beads (**Figure 1E**). These findings show that the LC can be reliably identified on the basis of its phase-translucence.

### Characterization of Neurons Acutely Dissociated from the LC

To identify the acutely dissociated LC cells as noradrenergic neurons, we used the immunocytochemistry and imaged the cells with widefield optics. Staining with a primary antibody against the neuronal dendritic marker MAP2 identified, as neurons, the large, multipolar cells present among those acutely dissociated from the phase-translucent LC brain sections (**Figure 2A,B**; MAP2). In addition, staining with an antibody raised against DBH revealed these neurons to be noradrenergic neurons (**Figure 2A,B**; DBH).

**Figure 2.**
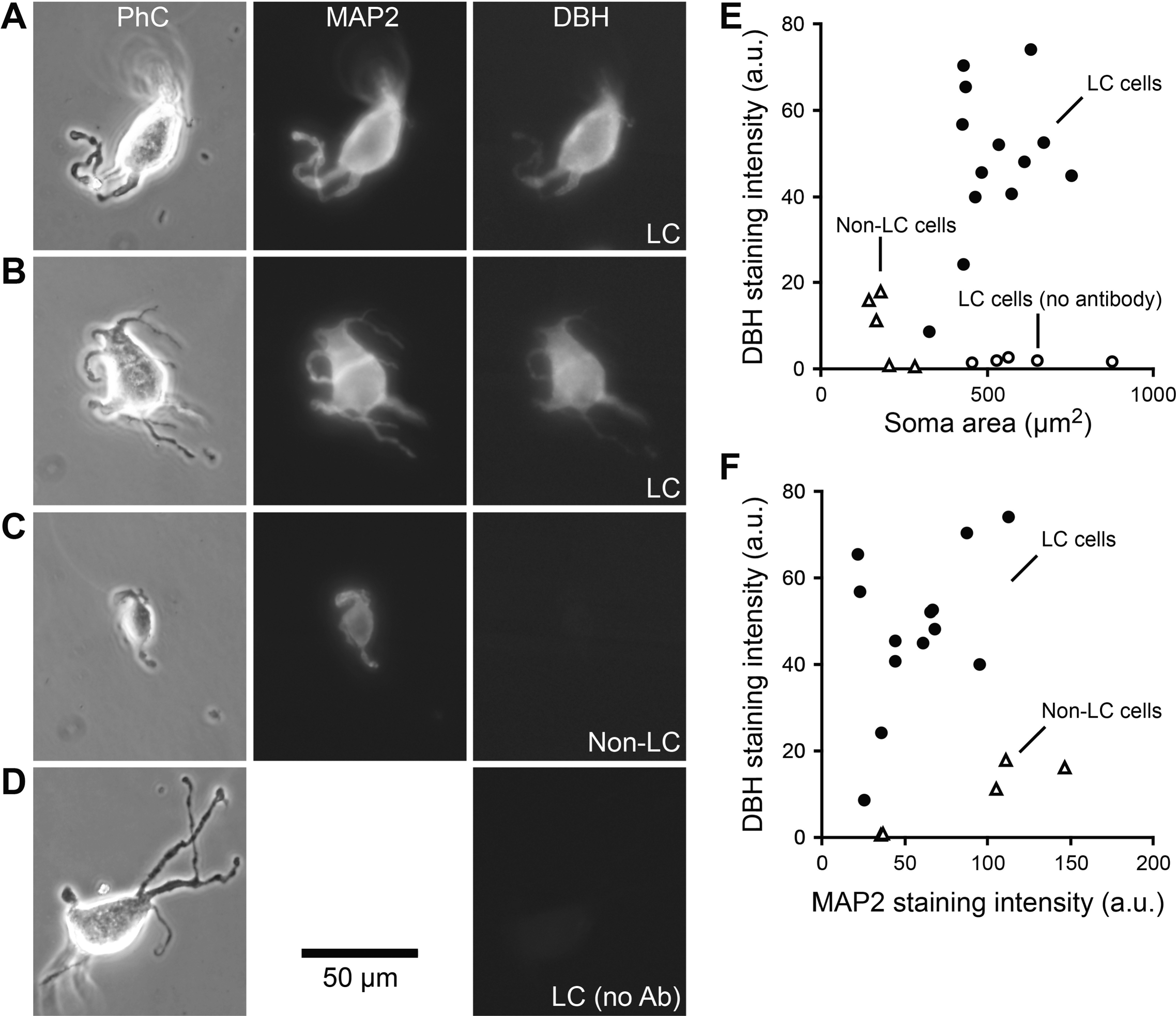
Identification of acutely dissociated cells from brainstem slices as noradrenergic neurons, using double-immunocytochemistry. **(A,B)** Two representative cells dissociated from the LC region (phase-contrast images) showed positive staining for the microtubule-associated protein 2 (MAP2, neuronal marker) and dopamine β-hydroxylase (DBH, noradrenergic marker). **(C)** Cells dissociated from a distinct region (non-LC, cochlear nucleus) were positive for MAP2 staining, but negative for DBH staining. **(D)** Negative control, in which the primary antibody against DBH was omitted. **(E)** Fluorescence intensity of DBH staining in individual cells, plotted against their somatic areas. Soma area was measured in phase-contrast image. **(F)** Fluorescence intensity of DBH staining plotted against that of MAP2 staining, for individual cells. Closed circles, LC cells; open triangles, non-LC cells (cells obtained from the cochlear nucleus); open circles, LC cells processed in the absence of primary antibody to DBH.

Two negative-control experiments were carried out to ascertain that our detection of DBH was specific for LC neurons. In one, we dissociated cells from the cochlear nucleus, a non-noradrenergic brainstem area. Cells acutely dissociated from this nucleus were positive for MAP2 but completely negative for DBH (**Figure 2C**). In another negative-control experiment, we omitted the primary, anti-DBH antibody from the immunocytochemistry procedure and failed to detect fluorescence labeling, showing that the fluorophore-conjugated secondary antibody is specific (**Figure 2D**).

Quantitative analysis of these data shows that DBH intensity in the soma of LC-derived neurons (47.7 ± 5.0 arbitrary units, a.u., n = 13 neurons, mean ± s.e.m.) was markedly stronger than that in either of the negative controls, i.e. neuronal soma derived from the cochlear nucleus (9.2 ± 3.7 a.u., n = 5 with p<1.0×10^−4^, unpaired t test), or neuronal soma derived from the LC but processed without primary antibody (1.7 ± 0.2 a.u., n = 5 with p<1.0×10^−6^) (**Figure 2E**). Further analysis revealed that the neurons isolated from the LC had considerably larger somatic area than their counterparts from the cochlear nucleus (523.0 ± 33.6 vs. 196.2 ± 23.9 µm^2^, n = 13 and 5, respectively, with p<1.0×10^−6^) (**Figure 2E**). However, the larger size of LC neurons alone does not explain their high DBH staining intensity: the MAP2 staining intensity in the soma was similar in cells dissociated from the LC and from the cochlear nucleus (58.0 ± 8.0 vs. 86.8 ± 21.9 a.u., n = 13 and 5, respectively, with p>0.2) (**Figure 2F**). Thus, the large, multipolar cells observed in our preparations were norepinephrine-containing LC neurons.

### Subcellular Distributions of MAP2 and DBH

Additionally we characterized the subcellular localization of the two markers. To this end, we used confocal microscopy which provides better spatial resolution than the widefield microscopy, especially in the z-(axial) direction. Immuno-reactivity of MAP2 was strong in dendrites, weak in the somatic cytoplasm, and absent in nucleus, as expected for a dendritic marker (two representative cells in **Figure 3A,C**). Background fluorescence was low, as indicated by a negative control from which the primary antibody for MAP2 was omitted (**Figure 3B**). The immuno-reactivity of DBH was absent from the nucleus, but in contrast to MAP2 staining, it was equally distributed throughout the cytoplasm in both dendrites and soma (six representative cells in **Figure 3D-H, J**). Background fluorescence was low, as indicated by a negative control from which the primary antibody for DBH was omitted (**Figure 3I**). These results were confirmed quantitatively by comparing the stained intensity of dendrites with respect to that of soma in the same cells (MAP2: p = 2.08×10^−12^, n = 32 ROIs in 13 cells; DBH: p = 0.44, n = 24 ROIs in 9 cells; paired t-test) (**Figure 3K**). These results indicate that the acutely dissociated cells show marker distributions as expected for intact LC neurons.

**Figure 3.**
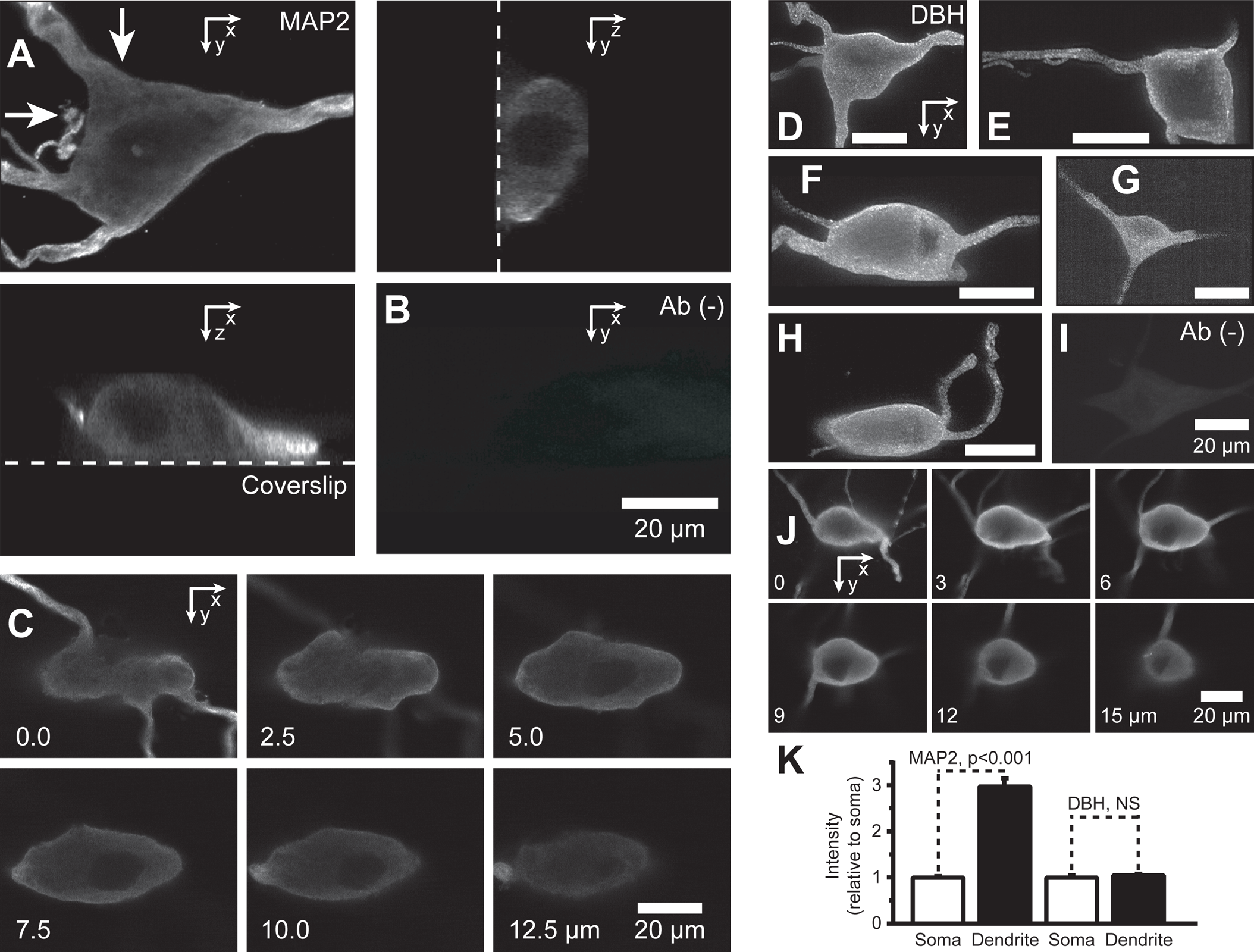
Subcellular Localizations of MAP2 and DBH. **(A)** Images of a MAP2-stained cell, visualized as: a maximum intensity projection of a stack with a horizontal view (x-y), a single vertical section (x-z), and another vertical section (y-z). The positions of the two vertical optical sections are indicated by arrows in x-y image (top left). Dotted lines indicate the positions of the coverslip. **(B)** Image of a cell in negative control (no primary antibody), visualized as a maximum intensity projection of a stack with a horizontal view. The scale bar in **(B)** applies to **(A)**. **(C)** Images of another MAP2-stained cell, visualized in individual horizontal sections. Numbers at bottom left indicate the height (in µm) of the x-y plane relative to the coverslip. **(D-H)** Maximum intensity projection of a z-stack image series taken from representative LC cells. All the scale bars indicate 20 μm. **(I)** Maximum intensity projection of a z-stack image series of a negative control LC cell (no primary antibody). **(J)** Individual, x-y horizontal sections of an LC cell. The numbers indicate the height (in µm) of the x-y plane relative to the coverslip. **(K)** Quantification of subcellular distribution of MAP2 and DBH in acutely dissociated LC cells. Fluorescence intensity in the dendritic cytoplasm was higher than that in the somatic cytoplasm in the case of MAP2 staining (p<0.001, n=32 regions of interest (ROIs), N=13 cells, paired *t*-test), but the ratio was similar for DBH staining (p>0.4, n=24 ROIs, N=9 cells, paired *t*-test). Error bars represent the SEM in all figures of this paper.

### Caffeine-Induced Membrane Potential and Current in LC Neurons

The response of the acutely dissociated LC neurons to caffeine was analyzed using the nystatin-perforated method of the patch-clamp technique. In a neuron recorded under the current-clamp mode, the resting membrane potential was −64 mV, and the neuron fired action potentials spontaneously (**Figure 4A**). The application of 10 mM caffeine rapidly induced membrane hyperpolarization, accompanied by a reduction in the firing of action potentials and an increase in membrane conductance. After caffeine washout, the membrane potential and firing pattern recovered to pre-application levels within 20-40 sec. This response was representative of 5 neurons, and is consistent with the notion that caffeine opens K^+^ channels on the plasma membrane.

**Figure 4.**
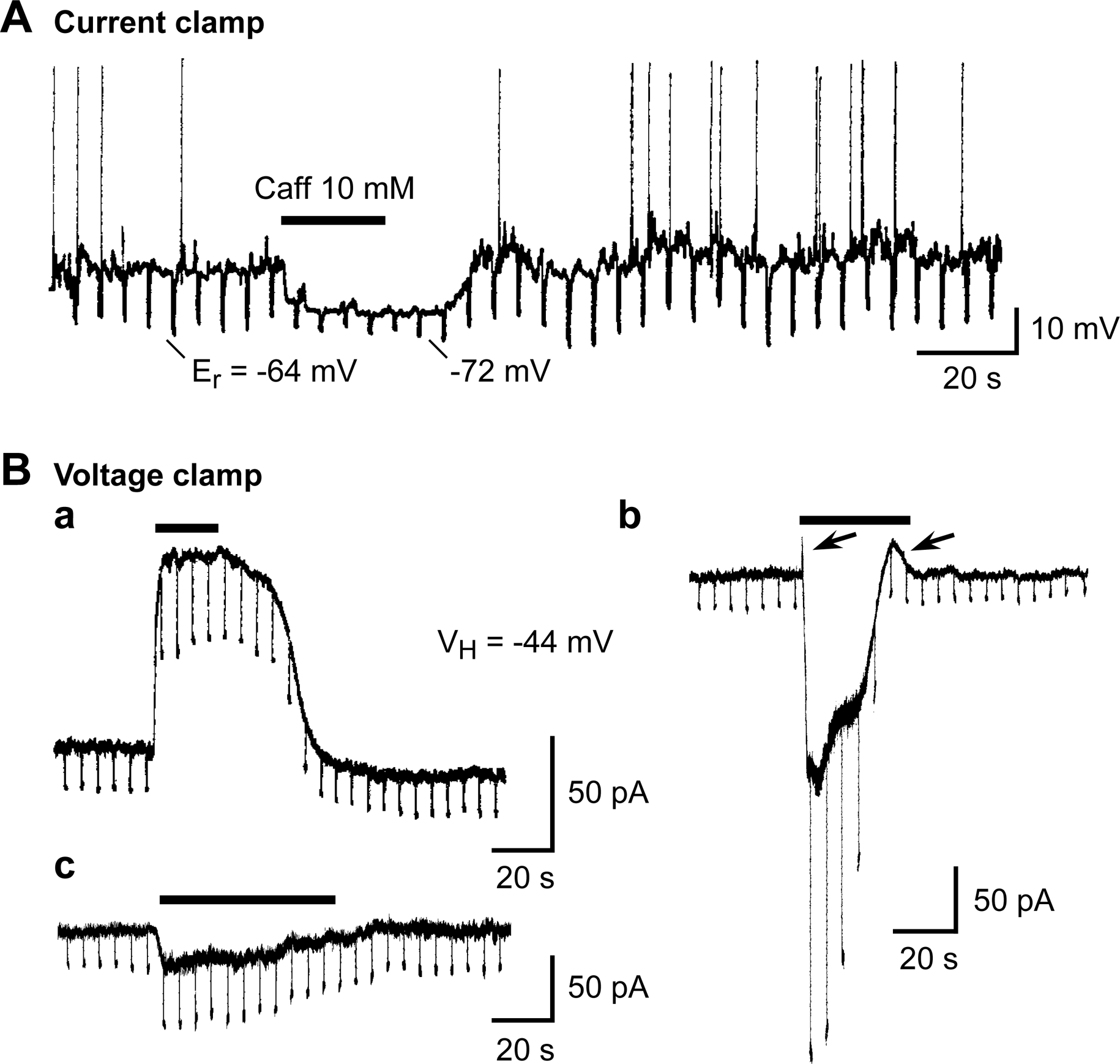
Electrophysiology of caffeine-induced effects in acutely dissociated LC neurons. **(A)** Current-clamp recording of responses to caffeine applied extracellularly (Caff, 10 mM). Several action potentials were recorded before caffeine was applied (large upward deflections). During caffeine application and slightly thereafter, the neuron was hyperpolarized (negative resting membrane potential, E_r_), and the firing of action potentials was suppressed. The membrane potential and action potential firing recovered to pre-application levels 20-40 sec after caffeine was washed out. Current pulses were repeatedly applied to the cell, at 5-sec intervals. A reduction in downward deflections during caffeine application indicates an increase in membrane conductance. **(B)** Voltage-clamp recording of the responses. Voltage pulses (−10 mV) were repeatedly applied to the cell, at 5-sec intervals. The increase in the amplitude of the downward deflections in all cells, represented by traces in **(Ba-c)**, supports an increase in membrane conductance. **(Ba)** In the majority of cells (78%), the application of 10 mM caffeine led only to an outward current. **(Bb)** In some cells (18%), three phases of response were observed: a small and transient outward current, a large inward current, and an outward current (the arrows point to outward currents). The overall membrane conductance increased over the course of treatment with caffeine. **(Bc)** In a small proportion of the cells (2%), the response consisted of a slowly recovering inward current accompanied by an increase in membrane conductance. Horizontal filled bars indicate the duration of caffeine application.

The caffeine response was analyzed more quantitatively using the voltage-clamp mode of the patch-clamp technique, at a holding potential of −44 mV. In the majority of neurons (39 of 50 neurons, 78%), caffeine application led to an outward current (**Figure 4Ba**), a finding compatible with the observation of membrane hyperpolarization in the current-clamp mode (**Figure 4A**). In the remaining neurons, caffeine induced one of three responses: one comprising an initial, transient outward current, a subsequent inward current, and a final outward current (9 of 50 neurons, 18%; **Figure 4Bb**); a second comprising an inward current followed by an outward current (1 of 50 neurons, 2%; data not shown); and a third characterized by an inward current only (1 of 50 neurons, 2%; **Figure 4Bc**). In all cases, the responses were accompanied by an increase in membrane conductance (as in the current-clamp measurements), implicating an opening of ion channels as the underlying mechanism.

### Kinetics of Currents Induced by 10 mM Caffeine

We reasoned that if these responses involve a multi-step process (diffusion of caffeine through the plasma membrane into the cytoplasm, sensitization of CICR, and a channel opening), there will be a delay in onset from the time of caffeine application, and there will be also certain variability in the detailed time courses. Thus, we examined the kinetics of caffeine-induced currents. These kinetics were compared with an internal standard of solution exchange in all recorded neurons. For this purpose, 30 mM K^+^ solution was applied (Kira et al., 1998), after the response to 10 mM caffeine has recovered, using the same drug application system. 30 mM K^+^ solution triggers an inward current by shifting the equilibrium potential of K^+^, and thus the time course of change in a holding current represents that of solution exchange in the cellular neighborhood. In all cases, the inward current induced by 30 mM K^+^ solution reached a steady-state level within 50 msec after onset (arrowheads in **Figures 5A-E**). This finding shows that the solution exchange was rapid, and that the concentration of the solution remained stable once the exchange was completed.

**Figure 5.**
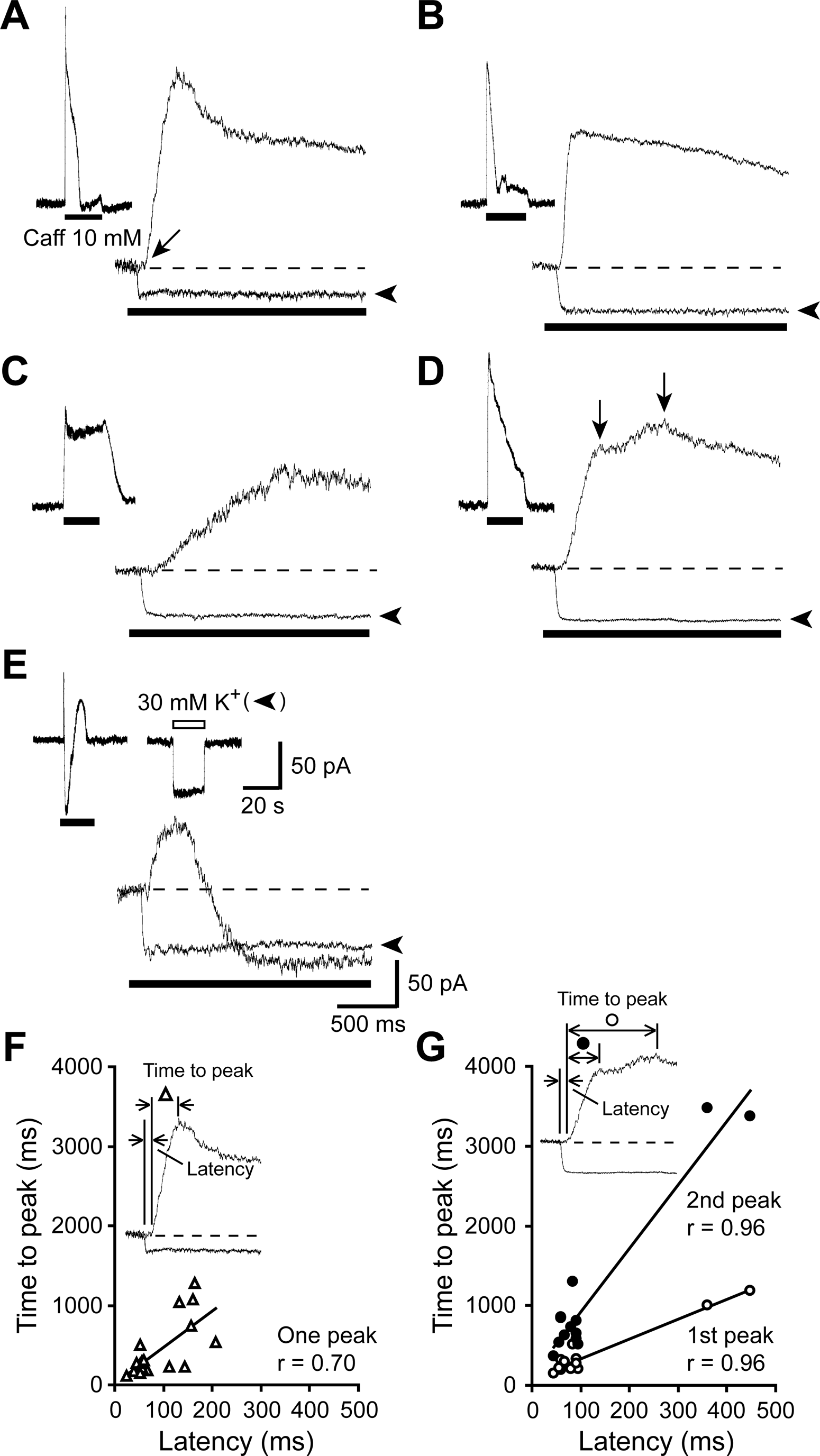
Variability in kinetics of the responses to caffeine. **(A-E)** Each panel shows a response of an acutely dissociated LC neuron to the application of 10 mM caffeine. Voltage-clamp recording was carried out at a holding potential of −44 mV. After the response to caffeine was recorded, 30 mM K^+^ solution was applied to the same neuron, to monitor the speed of solution exchange around the recoded neuron. The inset of each panel shows the response to caffeine on a slow time scale. The upper trace in the main part of each panel shows the response to caffeine on an expanded time scale, and the lower trace shows the response to 30 mM K^+^ in the same neuron and on the same time scale (arrowhead). Filled bars indicate the duration of caffeine or K^+^ solution application. Broken lines indicate the holding current before caffeine application. In panel **(A)**, an arrow indicates a later onset of caffeine response than the solution exchange. In panel **(D)**, vertical arrows indicate two peaks. Time scales and current scales for all the main panels and insets are indicated in panel **(E)**. One representative response to 30 mM K^+^ is shown at the slow time scale in the inset of panel **(E)**. **(F,G)** Relationship between the latency of onset and the time-to-peak for outward currents induced by 10 mM caffeine. Responses with a single peak **(F)** and two peaks **(G)**. The insets show how the parameters were measured. The latency was defined as the interval between 1) the time of initiation of solution exchange, as identified by the high K^+^-induced inward current, and 2) the time of onset of the outward current. The time to peak was defined as the interval between the onset of the outward current and the time at which the peak was reached. Pearson’s correlation coefficient (r) was 0.70 (p<0.001, n=19 neurons) **(F)**. The values were 0.96 (p<0.001, n=13 neurons **(G)**.

Kinetic analyses demonstrate several features of caffeine-induced currents. First, the caffeine-induced current always started much later than the solution exchange (**Figures 5F, G**). The latency was 103.6 ± 16.0 msec (n=32 cells), measured as the difference between the onsets of 30 mM K^+^-induced current and caffeine-induced current in each neuron (insets of **Figures 5F, G**). This long latency with respect to solution exchange is consistent with the idea that the observed current is produced by the multi-step process. Second, the caffeine-induced outward currents exhibited some variations: a transient peak followed by rapid decay (**Figure 5A**), a peak followed by a slow decay (**Figure 5B**), a slow rise to a peak with or without slow decay (**Figure 5C**), or two peaks (**Figure 5D**). These variations in the early phase of responses were often difficult to identify with slow time scales (insets). Third, a shorter latency was associated with a more rapid rise to peak. There was a positive correlation between the latency and the time to peak (measured as the time between the onset and the peak of caffeine-induced current; insets of **Figures 5F, G**). Pearson’s correlation coefficient (r) was 0.70 (p<0.001, n=19 neurons out of 32) for 1-peak cases (**Figure 5F**). The values were 0.96 (p<0.001, n=13 neurons out of 32) for both the first and the second peak of 2-peak cases (**Figure 5G**). Lastly, kinetic analysis revealed a notable feature with regard to the three-component response (equivalent to that shown in **Figure 4Bb**). Although the onset of the initial outward current (**Figure 5E**) was similar to that in other response types (**Figures 5A-D**), the outward current was abruptly replaced by an inward current approximately 400-500 msec after onset (**Figure 5E**). The late appearance of this inward current suggests that it is less sensitive than the initial outward current, to caffeine or its downstream consequences (such as increased [Ca^2+^]_c_).

### Dose-Response Relationship of Caffeine-Induced Outward Current

It is well established that caffeine sensitizes CICR in a narrow range of concentrations (from ∼1 to ∼10 mM) (Uneyama et al., 1993). Our analyses have similarly revealed that the outward current in rat LC neurons can be induced by caffeine at ∼3 mM (threshold level), and that maximal amplitudes are achieved at 10-30 mM (**Figure 6A**). This narrow and high concentration range in LC neurons is consistent with caffeine acting as a CICR-sensitizing agent.

**Figure 6.**
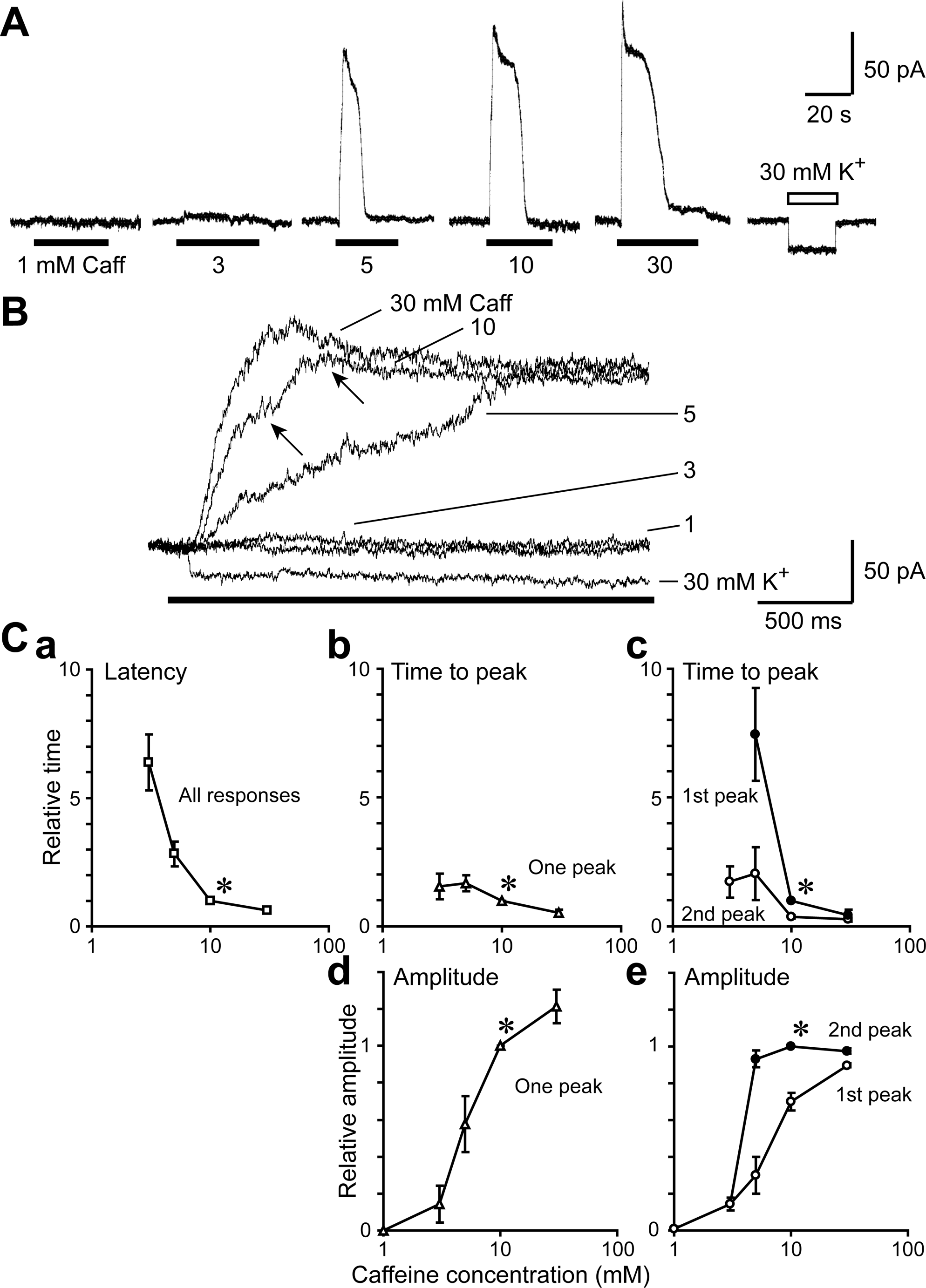
Dose-response relationship of caffeine-induced outward current in acutely dissociated LC neurons. Voltage-clamp recording was carried out at a holding potential of −44 mV. **(A)** Outward currents in response to different caffeine concentrations. 30 mM K^+^ (in the absence of caffeine) was applied at the end of the experiment to determine the speed of solution exchange. **(B)** Responses to different concentrations of caffeine by a single neuron are overlaid and presented on a faster time scale, to demonstrate the kinetics of current activation. In this cell, the outward current had two peaks (arrows). **(Ca)** Latency of responses at different caffeine concentrations, normalized to that at 10 mM caffeine (asterisk). **(Cb,c)** Time to peak of responses at different caffeine concentrations, normalized to that at 10 mM caffeine (asterisks). Responses with a single peak **(Cb)** and two peaks **(Cc)**. **(Cd,e)** Amplitudes of the outward current induced by caffeine at different concentrations. For each neuron, all amplitudes were normalized to that at 10 mM caffeine (asterisks). Responses with a single peak **(Cd)** and two peaks **(Ce)**. Each point represents the mean, and a vertical bar indicates ± s.e.m. where this exceeded the size of the symbol.

The kinetic analyses showed other, concentration-dependent features of the outward current (**Figure 6B**). With higher concentrations of caffeine, the current showed shorter latency (**Figure 6Ca**). This was accompanied by shorter time to peak for both 1-peak (**Figure 6Cb**) and 2-peak cases (**Figure 6Cc**). The dose-response relationship for the peak amplitudes showed similar concentration ranges (**Figure 6Cd,e**). These results clearly show that a concentration of caffeine is one factor that affects the latency and the time to peak of the outward current.

### Ca^2+^ Dependence of Caffeine-Induced Outward Currents within LC Neurons

The dose-response relationship suggests that the outward currents were induced by CICR which is not dependent on Ca^2+^ influx through the plasma membrane (**Figure 6**). If this is the case, the outward current should initially be insensitive to the removal of extracellular Ca^2+^, but gradually become sensitive to this condition, as the intracellular Ca^2+^ stores become depleted. The observed responses to caffeine were consistent with this notion. The response to 10 mM caffeine was unchanged when the cells were first incubated in the Ca^2+^-free extracellular solution containing 2 mM EGTA (Control & 1 min, **Figure 7A**). However, the response to a second application of caffeine in the Ca^2+^-free extracellular solution was less pronounced (6 min), and the response to a third application was negligible (11 min). This effect was reversible; the caffeine response was restored when the Ca^2+^-free extracellular solution was replaced with one at 2 mM Ca^2+^ (Recovery, **Figure 7A**).

**Figure 7.**
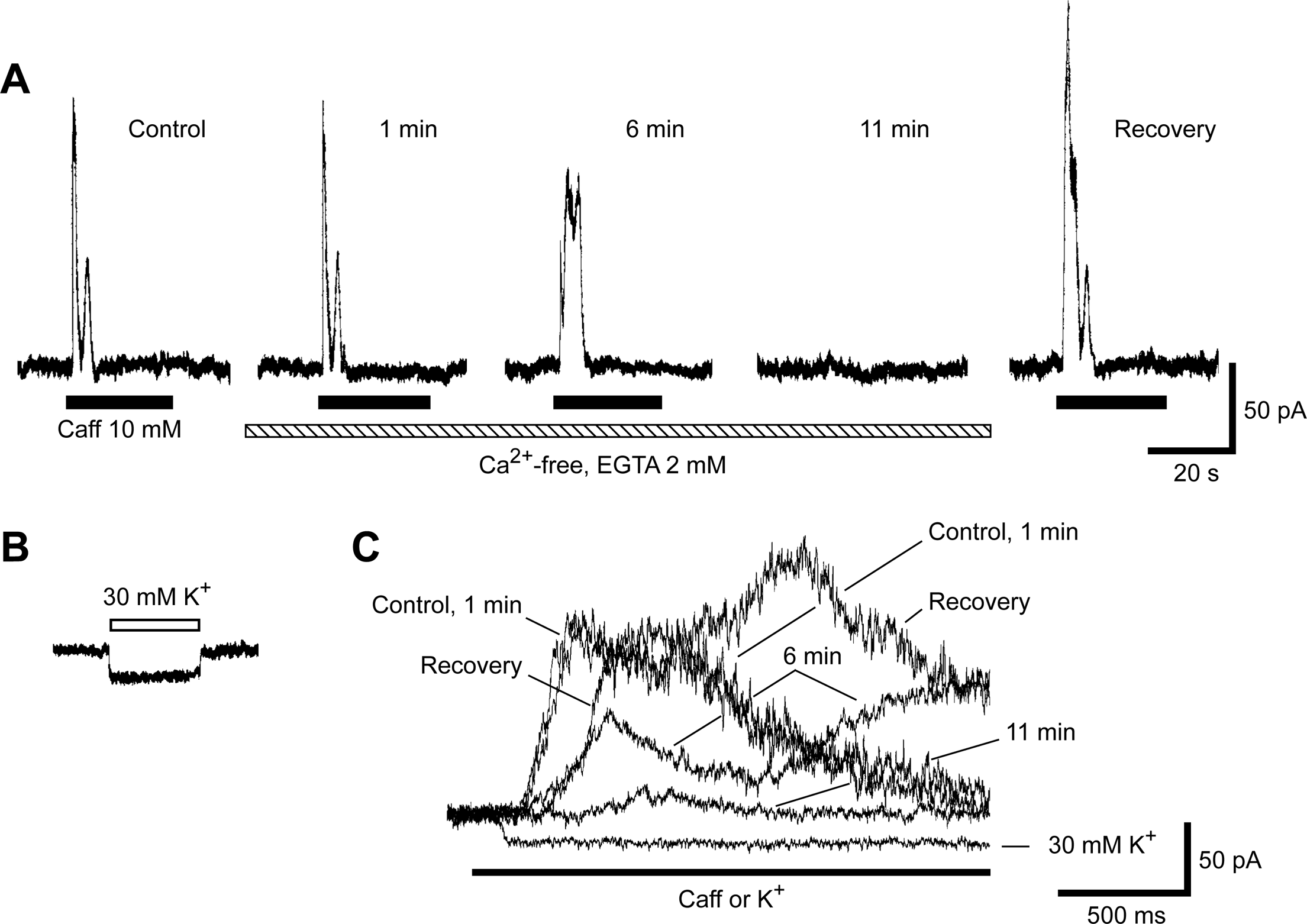
Ca^2+^-free extracellular solution abolished the caffeine-induced current gradually. Voltage-clamp recording was carried out at a holding potential of −44 mV. **(A)** The caffeine-induced outward current persisted for up to six minutes after the Ca^2+^-containing extracellular solution was replaced with Ca^2+^-free solution. Application of caffeine at 11 min elicited almost no response, indicating that the Ca^2+^ stores were depleted. Replacement of the external solution with Ca^2+^-containing solution restored the response to caffeine (recovery). **(B)** An inward current in response to 30 mM K^+^ solution in the same cell. **(C)** Overlay of responses to 10 mM caffeine and 30 mM K^+^, presented on an expanded time scale. These responses were representative of three neurons.

The kinetics of the responses were also analyzed in these neurons. **Figure 7B** shows the inward current in response to application of 30 mM K^+^ solution, used for assessing the solution exchange timing and efficiency (as in **Figures 5**,6). The caffeine-induced current rapidly rose to peak levels, before (control, **Figure 7C**) and at 1 min after application of Ca^2+^-free solution (1 min). However, after 6 min in Ca^2+^-free solution (6 min), the current exhibited slow rise to peak with two well-separated peaks. By the 11-min time point (11 min), the responses were suppressed to small amplitude with long latency and long time to peak. The response after replenishment of extracellular Ca^2+^ (Recovery) was accompanied by a relatively long latency, possibly reflecting incomplete recovery of Ca^2+^ loading into the intracellular stores.

Overall, these data suggest that the observed caffeine-induced outward current was not immediately dependent on extracellular Ca^2+^. They further indicate that the kinetics is also affected by the level of [Ca^2+^]_c_ or a degree of Ca^2+^ store filling that is reduced by continuous Ca^2+^-free extracellular solution (**Figure 7C**). It is estimated that this latter factor can also contribute to generating various responses to a fixed concentration of caffeine (**Figures 5A-D**).

### Blockade of Caffeine-Induced Currents by Ryanodine Application

The above results indicate that the intracellular Ca^2+^ stores are involved in the responses to caffeine. To demonstrate such an involvement more directly, we applied ryanodine, an agonist of ryanodine receptor, whose continuous activation leads to depletion of the intracellular Ca^2+^ stores. Application of 10 μM ryanodine resulted in gradual reduction of the amplitude of the caffeine-induced outward current (**Figure 8A**). This change was accompanied by a longer latency and slower rise to peak (**Figure 8C**), consistent with the idea that the degree of Ca^2+^ store filling is one determinant of kinetics.

**Figure 8.**
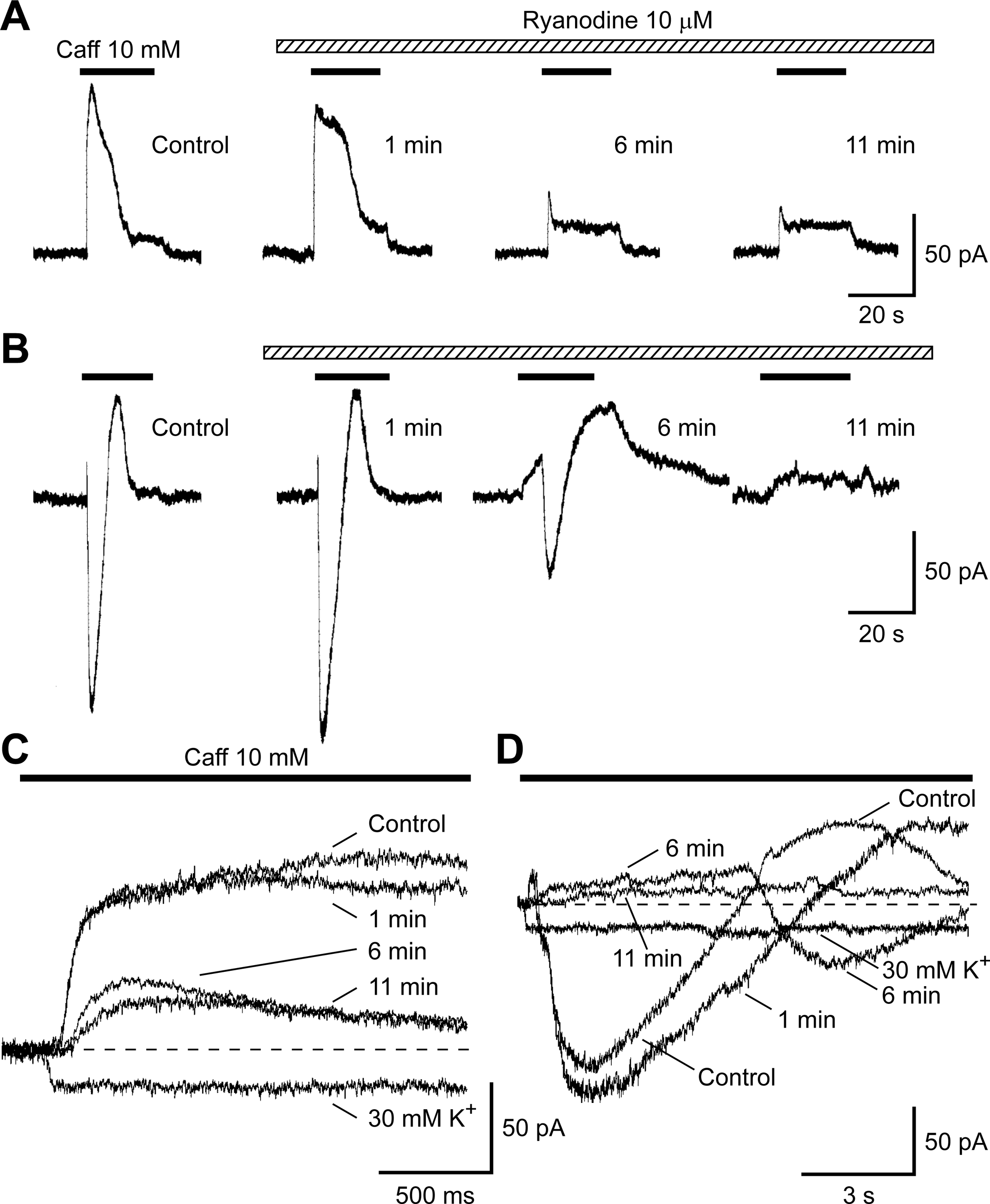
Sensitivity of the caffeine responses to Ca^2+^-store depletion by ryanodine. Voltage-clamp recording was carried out at a holding potential of −44 mV. **(A)** In neurons with a single-component response to caffeine, the amplitude of the outward current was gradually reduced during continuous treatment with 10 µM ryanodine. **(B)** Effects of ryanodine treatment, in a neuron with a three-component response to caffeine (initial outward, inward, and delayed outward currents). All components were abolished in parallel. **(C)** Overlay of the responses in panel **(A)**, presented on an expanded timescale. **(D)** Overlay of the responses shown in panel **(B)**, on an expanded timescale.

Also, in a cell exhibiting a three-component response to caffeine (consecutive outward, inward and outward currents), the ryanodine treatment led to parallel reductions in each component (**Figure 8B**). Interestingly, the onset of the inward current was slowed within 6 min of ryanodine treatment, as was the initial outward current (6 min, **Figure 8D**). These data suggest that both the outward and inward currents were mediated by Ca^2+^ release from intracellular Ca^2+^ stores.

### Current-Voltage Relationship of Caffeine-Induced Outward Current

Which ion channel is responsible for the caffeine-induced outward current? We approached this question by analyzing the current-voltage relationship (**Figure 9**). Ramp waves were generated by applying linear voltage ramps, both before and during exposure to 10 mM caffeine (a,b in inset of **Figure 9A**). The intersection of the two curves (**Figure 9Aa,b**) represents the reversal potential of the caffeine-induced outward current (E_Caff_). This was close to the calculated K^+^ equilibrium potential (E_K_) (**Figure 9A**). **Figure 9B** shows the effects of changing extracellular K^+^ concentration on E_K_, taking into account the effects of ionic activity (a[K^+^]_o_). A ten-fold change in a[K^+^]_o_ resulted in a shift of 56.8 mV for E_Caff_, in agreement with the predicted ∼59 mV at room temperature. These results indicate that K^+^ was the predominant charge carrier in producing the caffeine-induced outward current.

**Figure 9.**
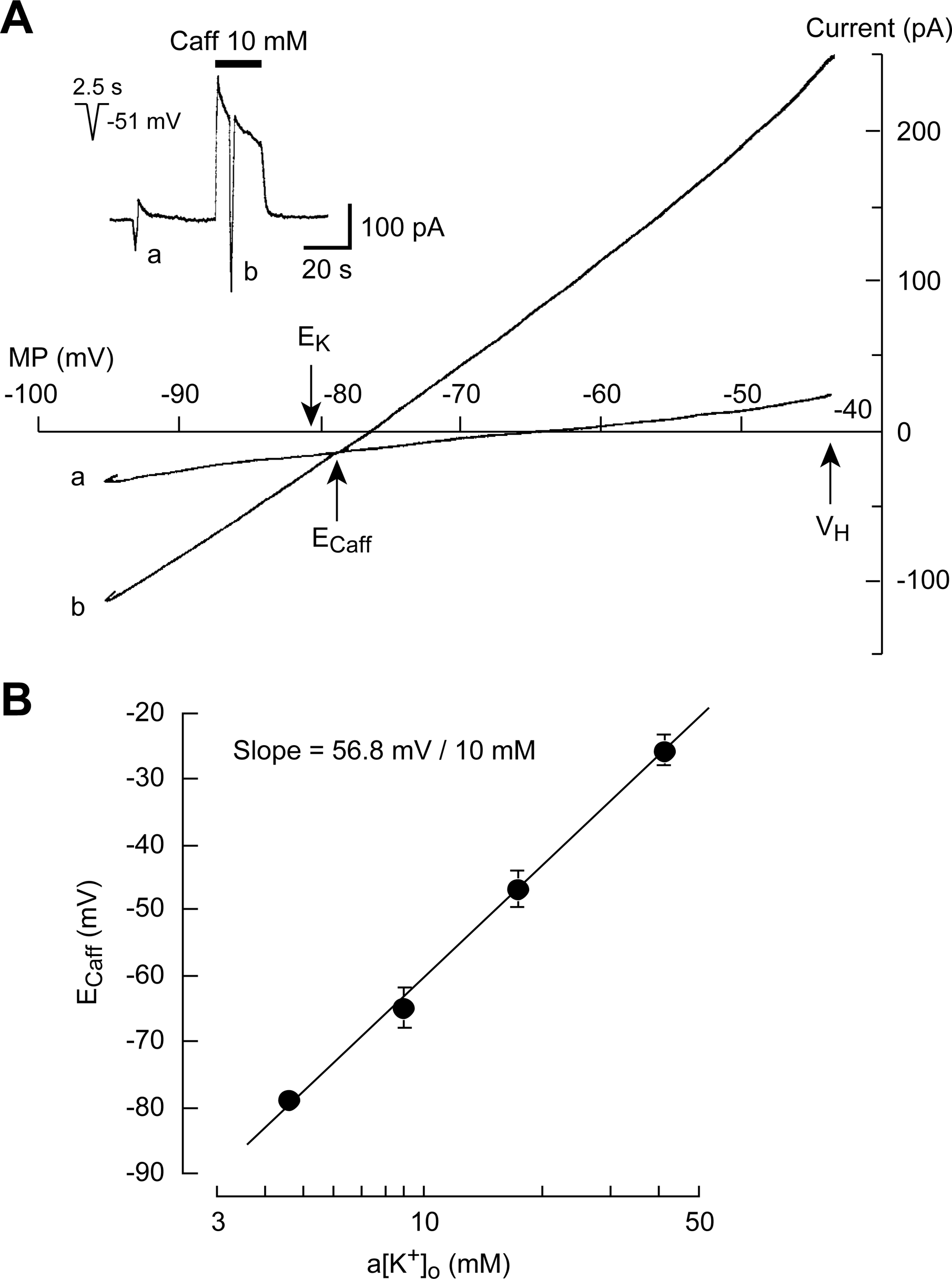
Current-voltage relationship of the caffeine-induced current. Voltage-clamp recording was carried out at a holding potential of −44 mV. **(A)** A voltage ramp was added to the holding membrane potential before **(a)** and during **(b)** the application of 10 mM caffeine. The inset shows the protocol for ramp-wave voltage command, with an amplitude of 51 mV and a total duration of 2.5 sec. The membrane currents in response to the voltage ramp crossed at a reversal potential (E_Caff_), which was near the K^+^ equilibrium potential (E_K_), as calculated based on the Nernst equation with the extra- and intracellular concentrations of K^+^. The extracellular K^+^ concentration, taking into account ionic activity (a[K^+^]_o_), was 4.6 mM. MP represents the membrane potential. **(B)** The reversal potential of the caffeine-induced current is plotted against different extracellular K^+^ activities. The E_Caff_ for a[K^+^]_o_ of 4.6, 9.0, 17.4 and 41.3 mM were −79.1 ± 0.97 (n = 4); −64.9 ± 2.98 (n = 3); −46.8 ± 2.81 (n = 3); −25.6 ± 2.40 (n = 3), respectively. The corresponding, calculated E_K_ values were −81.1, −64.3, −47.7 and −26.0 mV, respectively. A linear regression line had a slope of 56.8 mV per 10 mM change.

These results show that caffeine activates a K^+^ channel by sensitizing the release of Ca^2+^ from intracellular Ca^2+^ stores. Thus, the K^+^ channel responsible for the caffeine response is likely Ca^2+^-activated K^+^ channel. In order to characterize this channel, we assessed the effects of blockers of K^+^ channel types. The Ca^2+^-activated K^+^ channels are subdivided into small (SK), intermediate (IK) and large (BK) conductance types, which are blocked by apamin (SK), charybdotoxin (IK), and both iberiotoxin and charybdotoxin (BK). We pretreated the neurons with each of these inhibitors at a concentration of 0.3 μM for 1 min. Caffeine-induced outward current was not affected by iberiotoxin (Figure 10A), only partially inhibited by apamin (Figure 10A), and abolished by charybdotoxin (Figure 10B). These results suggest that caffeine opens mainly the Ca^2+^-activated K^+^ channels of the non-small conductance type.

**Figure 10.**
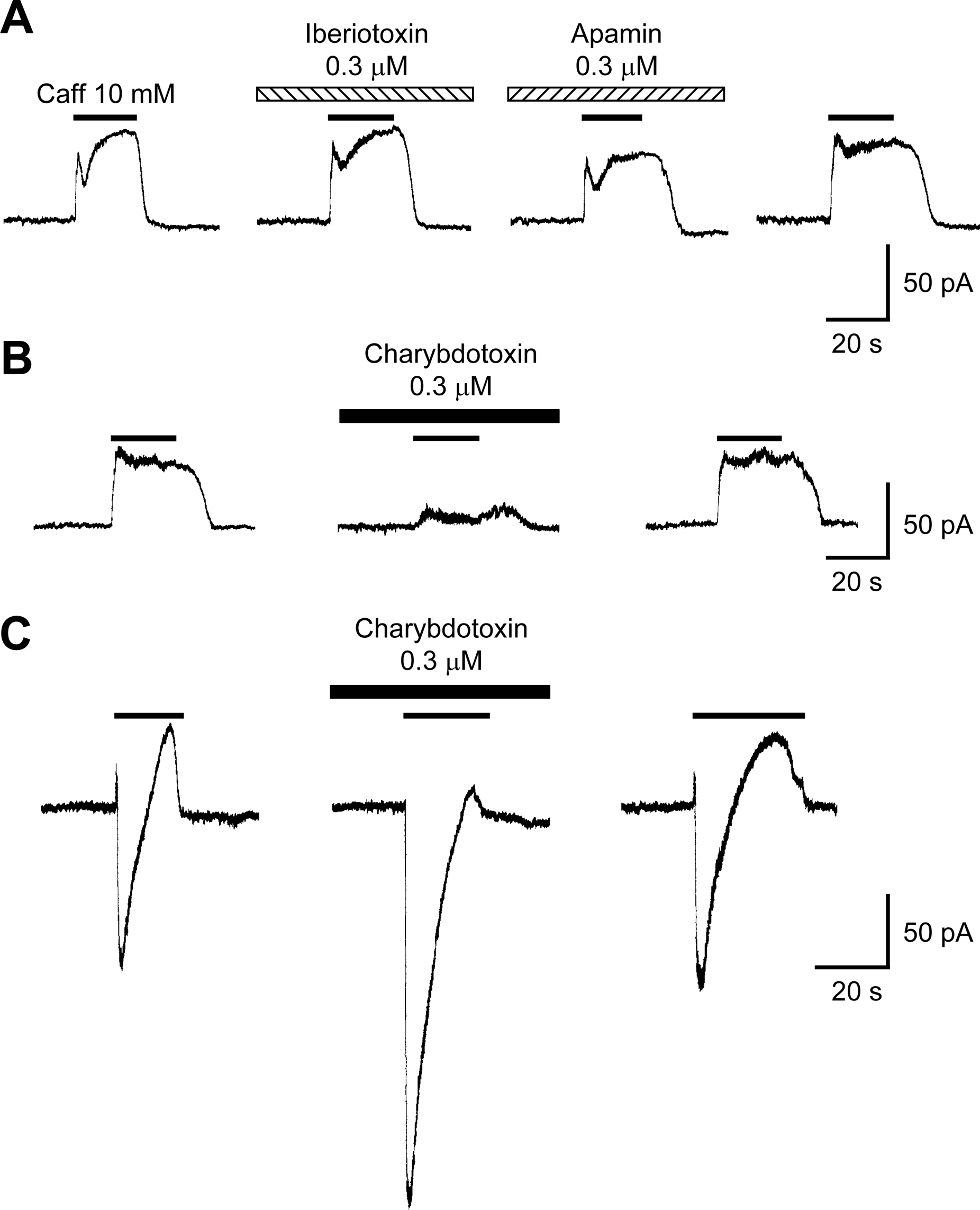
Pharmacology of K^+^ channels activated by caffeine. Voltage-clamp recording was carried out at a holding potential of −44 mV. **(A)** A neuron with a single-component response (outward current) to 10 mM caffeine. Neither treatment with 0.3 µM iberiotoxin, nor treatment with 0.3 µM apamin significantly affected the response to caffeine. **(B)** A neuron with a single-component response to 10 mM caffeine (similar to that in **A**). Treatment with 0.3 µM charybdotoxin resulted in a reversible block of the outward current. **(C)** A neuron with a three-component response to 10 mM caffeine. Treatment with 0.3 µM charybdotoxin blocked the outward current, unmasking a large inward current. This blockade was reversible.

In neurons that showed a three-component response to caffeine, the pretreatment with charybdotoxin likewise abolished the outward currents (Figure 10C). This blockade unmasked a large inward current. Therefore this response resulted from the sum of two distinct currents: 1) an outward, Ca^2+^-activated K^+^ current with early onset and long duration, and 2) an inward current with late onset and short duration. Unfortunately, the inward current could not be further analyzed due to its low incidence.

### Caffeine-Induced Ca^2+^ Mobilization and Its Dynamics

Finally we directly demonstrate that caffeine induces CICR, by imaging the changes in [Ca^2+^]_c_ based on a fluorescent Ca^2+^ indicator Fluo-3. We confirmed that a pretreatment with Fluo-3 did not affect the electrophysiological responses to caffeine (Supplementary Figure S2). When caffeine was applied continuously to the whole surface of LC neurons, the [Ca^2+^]_c_ increased in a manner consistent with caffeine-induced CICR from internal stores (Verkhratsky, 2005) (Figure 11Aa, with white arrows pointing in the direction of flow). This increase was observed throughout the cytoplasm and nucleus. However, there was subcellular variation in the kinetics of this increase (Figure 11Ab). The variation became more evident when the change in fluorescence intensity was normalized to the maximal value of each region (Figure 11Ac) and its slope was plotted (Figure 11Ad). Both representations indicate that the [Ca^2+^]_c_ increased more rapidly in a dendrite (red arrows) than in various somatic regions. The slowest response was observed at the center of soma, which corresponds to the nucleus (dark blue trace). This subcellular difference along the dendrite-to-soma axis was statistically significant (Figures 12A-C, p<0.05, p<0.001, p<0.001, respectively, Pearson’s correlation coefficient analysis, n=18 regions in 4 cells).

**Figure 11.**
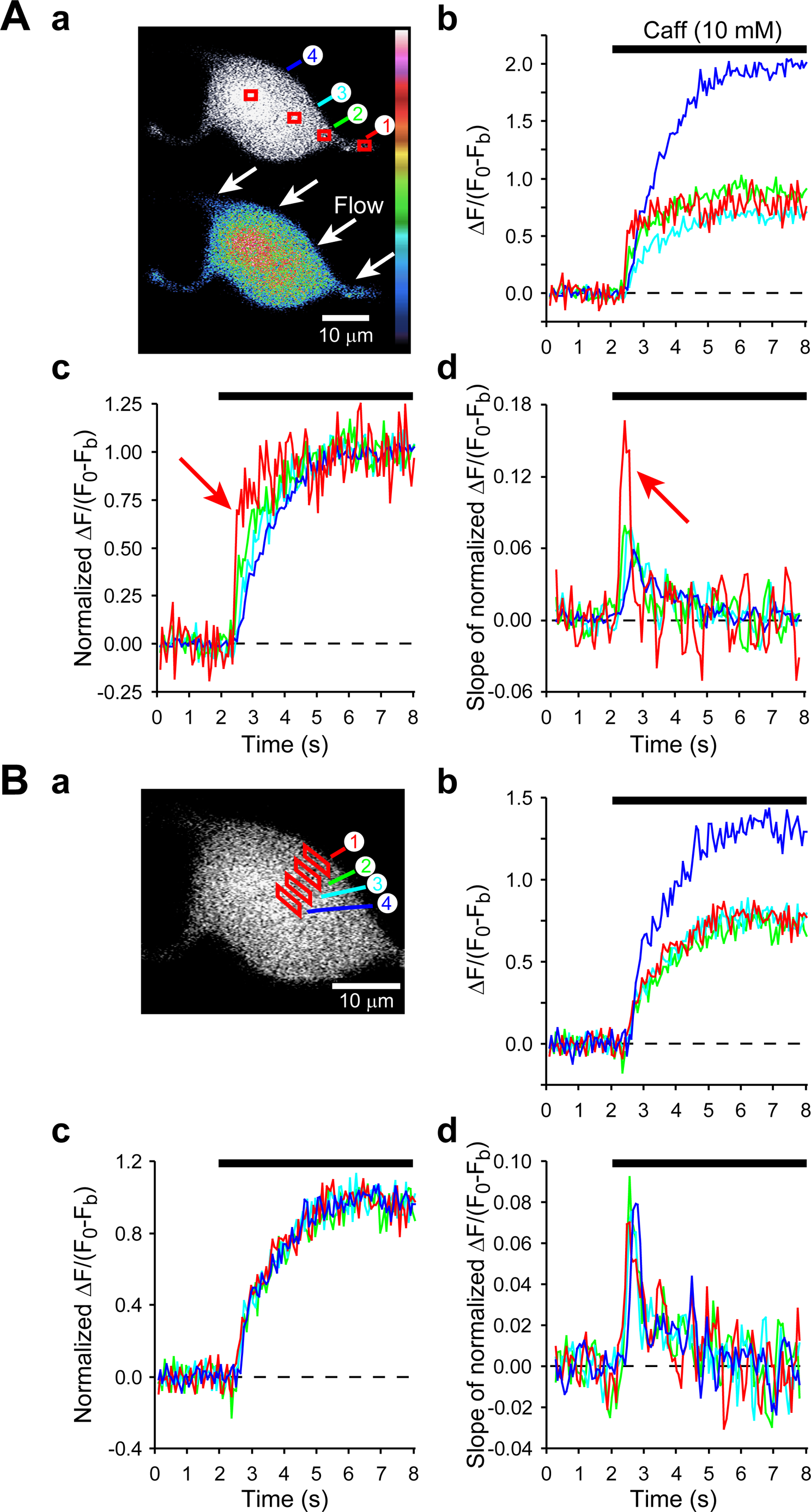
Caffeine (10 mM)-induced increases [Ca^2+^]_c_, as detected by confocal fluorescence imaging of Fluo-3-AM-treated LC neurons. Caffeine was applied to the whole surface of a dissociated LC neuron. The direction and timing of application are indicated by white arrows and black bars, respectively. **(A)** Analysis along the dendrite-to-soma axis. **(Aa)** Gray-scale (top) and pseudocolor-scale (bottom) images of a neuron responding to caffeine. **(Ab)** The changes in fluorescence intensity are plotted against time for ROIs shown in **(Aa)**. The colors of the ROI numbers and traces are matched. The change observed in the nuclear region of this cell (dark blue trace, ROI 4) was greater than those observed in a dendrite (red, ROI 1) and in other parts of soma (green, ROI 2; and cyan, ROI 3). **(Ac)** However, based on the normalized fluorescence change, the earliest and fastest rise was found in the dendrite (red trace, ROI 1, red arrow), and it became progressively later and slower with proximity to the nuclear region (dark blue trace, ROI 4). **(Ad)** The rate of rise in **(Ac)** was highest in the dendrite (red arrow). **(B)** Analysis along the surface-to-depth axis. **(Ba)** The cell in **(A)** was re-analyzed at different ROIs within the soma. **(Bb)** Fluorescence changes were larger in the nuclear region than in other parts of soma. However, normalized change **(Bc)** and the rate of change **(Bd)** were indistinguishable among ROIs at different depths. The Ca^2+^ signal is expressed as ΔF/(F_0_−F_b_), where ΔF is the change in fluorescence intensity from the pre-caffeine application intensity (F_0_), and F_b_ is the acellular background intensity outside the cells.

**Figure 12.**
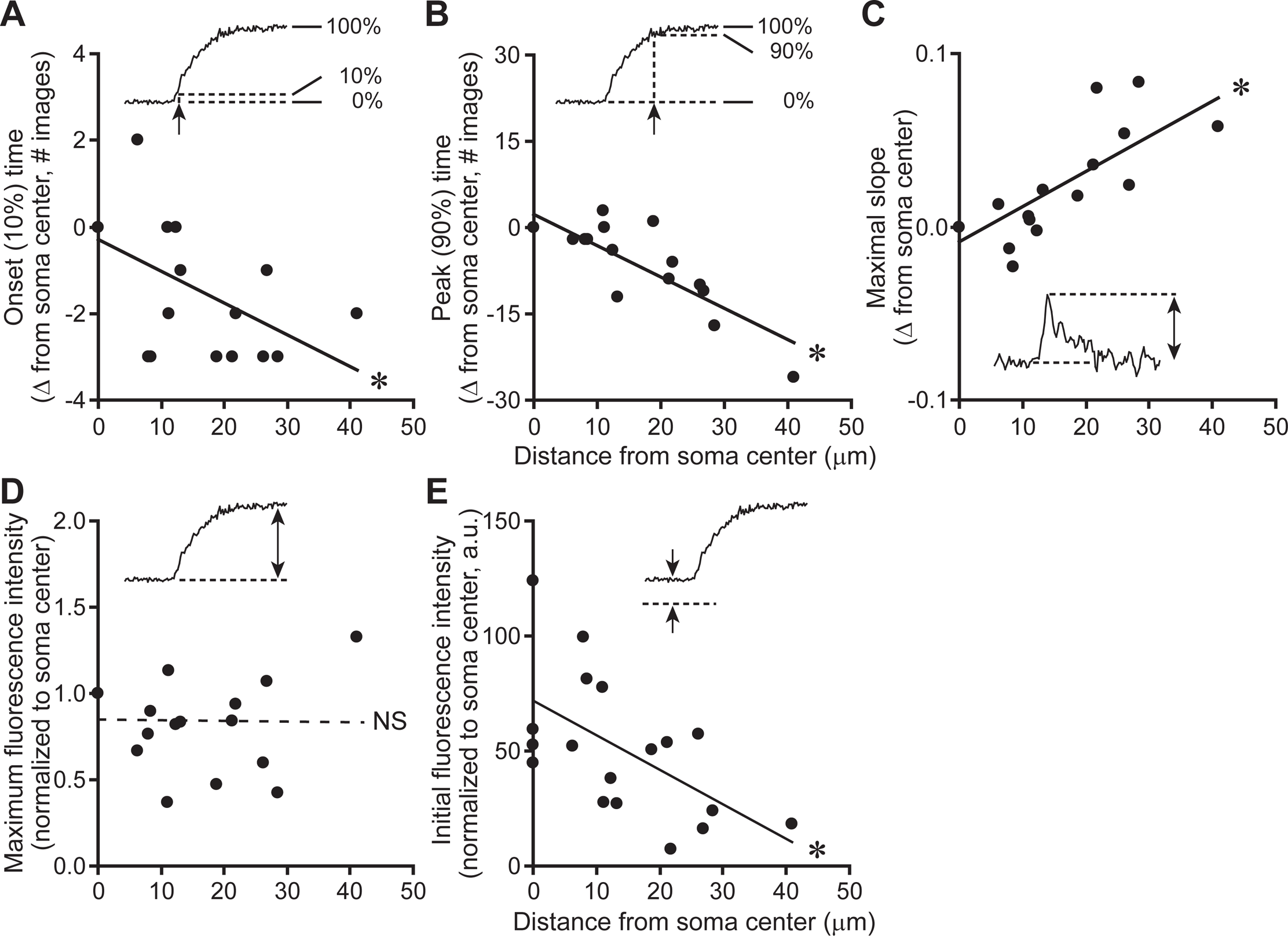
Quantitative comparisons of [Ca^2+^]_c_ increase dynamics, measured at various ROIs. All measured parameters are plotted against the distance of the ROI from the center of the soma. **(A)** The onset time of Ca^2+^ signal was earlier at more distant sites from the center of the soma (i.e. in the dendrites). Onset time was defined as the time when ΔF/(F_0_−F_b_) reached 10% of the maximum in each ROI. **(B)** The time of the Ca^2+^ signal peak was earlier at more distant sites from the center of soma (i.e. in the dendrites). Peak time was defined as the time when ΔF/(F_0_−F_b_) reached 90% of the maximum in each ROI. **(C)** The maximal slope of the Ca^2+^ signal was larger at more distant sites from the center of the soma (i.e. in the dendrites). The slope was calculated based on ΔF/(F_0_−F_b_), after peak intensity was normalized to 1. **(D)** Peak values of ΔF/(F_0_−F_b_) did not vary among the ROIs. **(E)** The initial, pre-caffeine intensity (F_0_−F_b_) was low in dendrites compared with that in soma. Statistical analysis was carried out using Pearson correlation analysis (n=18 ROIs in 4 cells).

Although it remains unclear why the kinetics of the fluorescence increase differed according to the regions, two possibilities can be excluded. First, it was not due to slow diffusion of caffeine from the plasma membrane toward deep cytosol. It could potentially have led to a later [Ca^2+^]_c_ rise in soma with a deeper cytosol. However, when the response in the same cell was analyzed at somatic ROIs at different depths along the surface-to-depth axis (Figure 11Ba), such subcellular differences were not noted (Figures 11Bb-d). This result indicates that our imaging rate (67 msec/frame) was too slow to detect either the caffeine diffusion within the cytosol, or if caffeine diffusion is slow, caffeine-triggered CICR in a superficial cytosolic layer rapidly propagating to deeper layers. In support of this notion, the speed of caffeine-induced Ca^2+^ wave was shown to be very rapid: in bullfrog sympathetic ganglion neurons, the Ca^2+^ wave propagated from the plasma membrane deeper into the cytosol at the speed of >44 μm/sec (Hua et al., 2000) (equivalent to >2.9 μm/67 msec or <3 frames for a neuron with ∼10 μm radius in our imaging study), which is below our temporal resolution. Second, the kinetic difference was not due to preferential saturation of Fluo-3 in dendrites, because there was no difference in the maximal fluorescence change in the dendrites *vs*. the somata (ΔF/(F_0_−F_b_), Figure 12D, p>0.1). The raw intensities of fluorescence before caffeine application were, in fact, smaller in dendrites than in somata (F_0_−F_b_, Figure 12E, p<0.05). We also found that the fluorescence signal decayed during constant caffeine application in some cells, and the decay was more rapid in the dendrites than in somatic regions (Supplementary Figure S3).

## DISCUSSION

This is the first systematic study of the effects of CICR activation on LC neurons. The overall scheme based on the current data in LC neurons was consistent with what has been reported for non-LC neurons (Supplementary Figure S4): caffeine application leads to the release of Ca^2+^ from the intracellular Ca^2+^ store into the cytosol, subsequent increase in [Ca^2+^]_c_, and opening of Ca^2+^-activated K^+^ channels. We also found some novel features in LC neurons. First, the sensitized CICR suppressed LC neuron excitability, mediated by channels whose pharmacology is compatible with the non-SK type. Second, there was heterogeneity in the kinetics of K^+^ current activation under different caffeine concentrations and during Ca^2+^ removal. Third, there was heterogeneity in the kinetics of caffeine-induced [Ca^2+^]_c_ increase and decrease in different parts of neurons.

### Morphological features of LC neurons

Although LC neurons have been acutely dissociated (Ingram et al., 1997; Murai et al., 1997; Nabekura et al., 1998; Chieng and Bekkers, 1999; Sengoku et al., 1999; Shirasaki et al., 2004; Koga et al., 2005; Ishibashi et al., 2009; Zhang et al., 2010), they have not been extensively characterized morphologically. In addition, identity of LC region has been questioned in some studies involving human brain imaging (Astafiev et al., 2010). Thus it was necessary to characterize the acutely dissociated LC cells. Our immunocytochemical data identified those cells that had been deemed live as neuronal (MAP2-positive) and noradrenergic (DBH-positive), the combination of which is the hallmark of LC neurons. The uniform distribution of DBH in the somato-dendritic region was consistent with the fact that LC neurons can release norepinephrine from the somato-dendritic regions (Huang et al., 2007).

### CICR in LC neurons

The [Ca^2+^]_c_-elevating action of caffeine in neurons was first proposed, based on the fact that an application of caffeine opened the Ca^2+^-activated K^+^ channels in bullfrog sympathetic ganglion neurons (Kuba, 1980). Ca^2+^ imaging was used to confirm that methylxanthines caffeine (Neering and McBurney, 1984) and theophylline (Smith et al., 1983) increase [Ca^2+^]_c_ by the CICR mechanism in vertebrate neurons. Since then, caffeine has been a compound of first choice in inducing the CICR-mediated increase in [Ca^2+^]_c_. In various types of neurons of the central and peripheral nervous systems, caffeine applied at 1 to 10 mM directly stimulates the CICR, increases [Ca^2+^]_c_, and induces an outward K^+^ current (Kuba, 1980; Marrion and Adams, 1992; Uneyama et al., 1993; Usachev et al., 1993; Garaschuk et al., 1997; Usachev and Thayer, 1997). Our electrophysiological and imaging data show that the same is true for LC neurons.

Previous electrophysiology studies had indicated that LC neurons support CICR. Specifically, it had been shown that the afterhyperpolarization following action potentials was mediated by two components of Ca^2+^-activated K^+^ currents (Aghajanian et al., 1983; Matschke et al., 2018), and that the slow component was suppressed by treatment with ryanodine (known to block the CICR) (Osmanovic and Shefner, 1993). Our Ca^2+^ imaging analysis extends those indirect findings in LC neurons, by clearly demonstrating that the ClCR leads to an increase in [Ca^2+^]_c_.

### Properties of caffeine-induced outward current

The basic properties of the caffeine-induced currents in LC neurons were similar to those of currents previously observed in non-LC neurons, in terms of: effective concentration of caffeine, sensitivity to removal of extracellular Ca^2+^, sensitivity to ryanodine, and activation of K^+^ channels (Marrion and Adams, 1992; Uneyama et al., 1993; Garaschuk et al., 1997; Coulon et al., 2009). Additionally, our kinetic analyses of the latency and the rise to peak provide new information about at least two regulating factors of caffeine-induced currents in LC neurons. One is the caffeine concentration. The latency and the time to peak were negatively correlated with the caffeine concentration (**Figure 6**), which will affect the sensitivity of CICR to Ca^2+^ in the multi-step process involving permeation through the plasma membrane, diffusion of caffeine in the cytoplasm, sensitization of CICR, and a channel opening. Another factor is the extent to which Ca^2+^ level is maintained. The latency and the rise to peak were prolonged when the intracellular Ca^2+^ stores were depleted by removing extracellular Ca^2+^ (**Figure 7**) or treating with ryanodine (**Figure 8**). These findings indicate that the efficiency of CICR is affected by the levels of [Ca^2+^]_c_ and/or Ca^2+^ store filling, consistent with findings in sympathetic neurons (Hernandez-Cruz et al., 1997; Albrecht et al., 2001). This interpretation further indicates that variation in the latency and the time to peak among neurons (**Figure 5**) partly reflects natural variation in Ca^2+^ levels at the resting state.

### Identity of Ca^2+^-activated K^+^ channels

Ca^2+^-activated K^+^ channels show different sensitivities to pharmacological inhibitors. SK is sensitive to a bee-venom apamin, with the half-maximal inhibitory concentration (IC_50_) of 0.83-3.3 nM (Stocker, 2004; Honrath et al., 2017; Matschke et al., 2018). IK is sensitive to scorpion toxin charybdotoxin (Stocker, 2004; Honrath et al., 2017) with an IC_50_ of 2-28 nM (Pedarzani and Stocker, 2008), and insensitive to apamin (Pedarzani and Stocker, 2008; Engbers et al., 2012). BK is in general sensitive to scorpion toxins charybdotoxin with an IC_50_ of 3 nM (Honrath et al., 2017), and iberiotoxin with an IC_50_ of 0.36-0.76 nM (Salkoff et al., 2006; Honrath et al., 2017). However, some forms of BK (ones that contain β4 subunit) are resistant to both toxins (Meera et al., 2000; Salkoff et al., 2006; Wang et al., 2014; Gonzalez-Perez and Lingle, 2019). BK channels are insensitive to an SK channel blocker apamin.

The caffeine-induced Ca^2+^-activated K^+^ current assessed here was sensitive to charybdotoxin, but insensitive to iberiotoxin and apamin (**Figure 10**), and thus appears to be compatible with non-SK channels, especially the IK channels. A Ca^2+^-activated K^+^ current with similar pharmacological properties was observed in acutely dissociated LC neurons when the cells were subjected to ischemic conditions (Murai et al., 1997). In support of these data, the IK mRNA (*Kcnn4*) has been detected at a moderate level in the LC region of mouse brain (Allen_Mouse_Brain_Atlas, 2004) (image numbers 45 and 46 out of 56).

Why was the SK current absent in our experimental system? SK current was responsible for the fast phase of after-hyperpolarization in LC neurons within brain slices, as demonstrated by its sensitivity to apamin (Osmanovic et al., 1990; Osmanovic and Shefner, 1993; Zhang et al., 2010; Matschke et al., 2018). It thus seemed reasonable to expect that SK currents are activated by CICR in our system. Either of two possibilities could account for the lack of SK as a major effector. One possibility is related to the subtype and subcellular distribution of SK. Among three SK subtypes (SK1, SK2, SK3), the predominant form in LC neurons is SK3 (*KCNN3*), both in terms of mRNA (Stocker and Pedarzani, 2000) and protein (Sailer et al., 2004) expression levels. However, the bulk of its protein is present in varicose fibers (Sailer et al., 2004), which are absent from our acutely dissociated preparation. A second possible explanation for our results is that CICR may not be coupled to SK channels. Indeed, the coupling between CICR and particular Ca^2+^-activated K^+^ channel seems to be highly dependent on cell type. For example, CICR is coupled with SK currents, but not BK currents, in neurons from the rat superior cervical ganglion (Davies et al., 1996), the rabbit vagal nodose ganglion (Cordoba-Rodriguez et al., 1999) and the rat thalamic reticular nucleus (Coulon et al., 2009). Conversely, CICR is strongly coupled with BK currents, but only weakly with SK currents, in the mouse cartwheel inhibitory interneurons of the dorsal cochlear nucleus (Irie and Trussell, 2017; Irie, 2019) and in bullfrog sympathetic ganglion cells (Akita and Kuba, 2000). Thus, it is possible that CICR is coupled preferentially with non-SK channels in LC neurons.

### Caffeine-induced inward current

Caffeine induced an inward current in a small number of LC neurons. The inward current was associated with an increase in membrane conductance (**Figures 4Bb, c**), indicating an opening of ion channels. Ryanodine suppressed the amplitude and slowed the kinetics of the inward current, similarly as those on the outward current (**Figure 8**). This result strongly suggests that the two currents share the same activation mechanism. These results indicate that the inward current was activated by opening of ion channels as a consequence of CICR. The later onset of the inward *vs.* outward current (**Figures 5E**,8D) suggests that this Ca^2+^-activated ion channel has a lower affinity for cytosolic Ca^2+^ than does the K^+^ channel that mediates the outward current. Studies in non-LC neurons have identified at least two types of ionic current that could potentially account for the inward current observed here. One is a Ca^2+^-activated Cl^-^ current found in sympathetic neurons of the bull-frog (Akaike and Sadoshima, 1989) and mouse (Martinez-Pinna et al., 2000), dorsal root ganglion neurons of the rat (Currie and Scott, 1992; Ayar and Scott, 1999) and chick (Ward and Kenyon, 2000), and vagal nodose neurons of the rat after vagotomy (Lancaster et al., 2002). Another is a Ca^2+^-activated non-selective cation current, found, e.g., in dorsal root ganglion neurons of rat (Ayar and Scott, 1999). The identification of this current in LC neurons will need a further study.

### Caffeine-induced [Ca^2+^]_c_ kinetics in LC neurons

We found subcellular differences in the kinetics of CICR-induced [Ca^2+^]_c_ changes. The rising and decay phases of [Ca^2+^]_c_ were more rapid in dendrites than in somata, even though caffeine was applied continuously throughout the neuronal surface. These differences in CICR kinetics in the dendrite-to-soma direction are novel. Both of these subcellular differences can be most easily explained by the different surface-to-volume ratios in dendrites and soma. The surface-to-volume ratio represents the surface area of plasma membrane per unit volume of the structure, i.e. the relative abundance of plasma membrane with respect to the sum of cytosol and organelles. This value is higher for dendrites because of the smaller diameter than the soma. The surface-to-volume ratio in dendrites has been discussed for interpreting the action potential-induced [Ca^2+^]_c_ changes, in terms of larger amplitudes in cerebral cortical pyramidal neurons (Cornelisse et al., 2007) and faster decay (Lev-Ram et al., 1992) in cerebellar Purkinje neurons.

In the current experimental system, the degree of caffeine access to the cell is proportional to the surface area, and thus the effect of caffeine is expected to be larger in dendrites, assuming that the Ca^2+^ store density is similar throughout a neuron. This could explain the earlier and faster rise kinetics in dendrites in LC neurons. The [Ca^2+^]_c_ was reported to decrease in the continued presence of caffeine (Usachev et al., 1993; Choi et al., 2006). Activity of plasma membrane Ca^2+^-ATPase (PMCA), a Ca^2+^ pump that transports Ca^2+^ out of the cytosol to the extracellular space, was partly responsible for the [Ca^2+^]_c_ decay during caffeine application (Usachev et al., 1993) and after action potentials (Usachev et al., 2002) in the soma of rat dorsal root ganglion neurons. This will lead to more effective Ca^2+^ extrusion and therefore more rapid [Ca^2+^]_c_ decay in dendrites than in somata. The elucidation of these mechanisms will require detailed analysis of the factors known to affect [Ca^2+^]_c_, such as the regional densities of PMCA, Na^+^/Ca^2+^ exchanger on the plasma membrane, Ca^2+^-binding proteins in the cytosol, RyR and the endoplasmic reticulum as the Ca^2+^ pools (Berridge et al., 2003; Friel and Chiel, 2008; Schwaller, 2009).

### Roles of CICR in LC neurons

It seems that all neuronal types that have been examined to date undergo CICR (Verkhratsky, 2005), including LC neurons (this study). However, this is in sharp contrast to some other cell types that vary in their ability to support CICR. For example, smooth-muscle tissue is very heterogeneous: CICR occurs frequently in the guinea-pig ureter (Burdyga et al., 1995), occurs in 40% of guinea-pig taenia caeci (Iino, 1990), occurs in 30% of the rat myometrium (Martin et al., 1999), and fails to occur in the rat ureter (Burdyga et al., 1995). CICR is not well developed in astrocytes compared with that in central neurons (Beck et al., 2004). The apparently universal occurrence of CICR in neurons, in contrast, implies that it plays an essential role in neuronal functions.

One of the major roles of CICR in LC neurons will be to suppress membrane excitability (**Figure 4A**), through triggering Ca^2+^-activated K^+^ currents that contribute to excitability control by regulating firing frequency and spike adaptation (Osmanovic and Shefner, 1993; Stocker, 2004). Additionally, CICR could regulate LC neuronal excitability by other means as well, for example by limiting the spatial and temporal ranges of dendritic Ca^2+^ spikes, as is the case for BK-type Ca^2+^-activated K^+^ currents in cerebellar Purkinje neurons (Rancz and Hausser, 2006), and SK-type currents in hippocampal pyramidal neurons (Cai et al., 2004). CICR is also involved in the regulation of synaptic physiology, such as in hippocampal dendritic spines (Emptage et al., 1999), nerve terminals (Emptage et al., 2001), cortico-striatal synaptic plasticity (Popescu et al., 2010), and in pathophysiology, such as aging process (Gant et al., 2006) and Alzheimer’s disease (Goussakov et al., 2010; Popugaeva et al., 2017). Thus CICR is likely to be one of the essential means of regulating the LC neuronal activity. Based on the innervation of widespread areas of the CNS by even a single neuron (Foote et al., 1983; Berridge and Waterhouse, 2003; Schwarz et al., 2015), LC neuron can influence the noradrenergic tone throughout the nervous system. CICR in LC neurons can be involved in pathological conditions, e.g. autonomic dysfunction as in neurogenic orthostatic hypotension.

## AUTHOR CONTRIBUTIONS

Conception and design of the study: NCH. Acquisition, analysis and interpretation of data: HK, SBM, JYK, KMG, NCH. Drafting the manuscript: NCH, and all the authors critically revised the draft and approved the final version.

## ACKNOWLEDGMENTS

The authors thank Dr. Norio Akaike for a generous access to the Noran confocal microscope.

## SUPPLEMENTARY MATERIALS AND METHODS

### Fluorescent, live/dead staining

The Live/Dead Fixable Red Dead Cell Stain kit was used (L23102, Molecular Probes-Invitrogen, Carlsbad, California, USA). One vial was reconstituted in 50 μl of dimethylsulfoxide (DMSO) according to the manufacturer’s instructions, and further diluted 1000-fold in the standard external solution. The acutely dissociated cells were treated with this dye for 30 min at room temperature, and rinsed with dye-free standard external solution 4 times for 1 min each. The cells were observed under an widefield fluorescence microscope (Eclipse TE300, Nikon) using a light source (X-Cite 120, EXFO Life Sciences Division, Mississauga, Ontario, Canada), a fluorescence filter cube (exciter: HQ545/30; emitter: HQ610/75; beam splitter: Q570LP) and 40x air objective lens (Plan Fluor ELWD, NA 0.60). The images were acquired using a CCD camera (DP71, Olympus, Center Valley, Pennsylvania, USA), with an exposure time of 2 sec. The fluorescence intensity of a cell was measured by defining a region of interest (ROI) around the perimeter of the soma (identified in the phase-contrast image) and averaging the pixel intensity within the ROI using ImageJ. The intensity was corrected by subtracting the fluorescence intensity outside the cells (acellular background).

## SUPPLEMENTARY RESULTS / DISCUSSION

### Identification of Acutely Dissociated LC Cells as Live Neurons

In all experiments in the current report (live-cell electrophysiology, live-cell [Ca^2+^]c imaging, and fixed-cell immunocytochemistry), the acutely dissociated cells need to be alive (or alive before chemical fixation), and therefore free of pathological modulations. This is important because our result will be affected if the [Ca^2+^]c rises, for example in the context of ischemia (Silver and Erecinska, 1990; Murai et al., 1997). It is also important that the cells are alive immediately before chemical fixation for immunocytochemical analyses, because intracellular proteins are known to be redistributed under cellular stresses, and their normal subcellular distribution would be masked. For example, MAP2 is normally abundant in the dendrites of pyramidal neurons (Craig and Banker, 1994), whereas it will be lost from dendrites and redistributed throughout the somata under ischemia and excitotoxicity (Hoskison et al., 2007).

It is common practice for investigators to determine the live (healthy) condition of the cells by imaging with transmitted light microscopy (e.g. phase-contrast and differential interference contrast microscopy). We confirmed the validity of such transmitted light-based approach, by using fluorescent, live/dead cell staining, and comparing the results obtained using the two methods. The live/dead assay is based on the fact that live cells expel a fluorescent dye, and therefore exhibit low fluorescence intensity (Figure S1A), whereas dead cells (whose plasma membrane is damaged) are permeated by the fluorophore and exhibit high fluorescence intensity (Figure S1B). In one representative dissociation series, 11 cells were subjected to live/dead staining and to a blind test of classification. Two experienced experimenters observed only the phase-contrast images and decided whether the cells were alive or dead (see Materials and Methods for criteria). Later, the matching images of the fluorescence from the live/dead staining were used to assess the condition of each cell. The live/dead staining indeed led to different fluorescent intensities of cells (p<0.001, *t*-test) (Figure S1C). The cells determined to be alive by two experimenters based on phase-contrast microscopy showed lower fluorescence intensity following live/dead staining than the cells determined to be dead (p<0.005 and p<0.02, Fisher’s exact probability test, Figure S1D). The fluorescence intensity was not positively correlated with the area of soma (p>0.1, n=11 cells, data not shown), indicating that cell size is not a major determinant of cell viability. These results confirm that the evaluation of transmitted-light images is an effective means of determining cell viability.

The evaluation of the transmitted-light images is also useful even after chemical fixation with paraformaldehyde, because chemical fixation did not change the phase-contrast properties of acutely dissociated cells (data not shown). Thus, for obtaining images of immunocytochemically stained cells, we first observed the phase-contrast images of the fixed cells, and used only those cells with “live-cell” features for acquiring fixed-cell images.

The live condition of the acutely dissociated cells was further tested using the patch-clamp electrophysiological recording. An inward current was induced in the voltage-clamp mode, when the membrane potential was held at –80 mV and stepped to 0 mV (Figure S1E). The induced inward current was completely blocked by 1 µM tetrodotoxin, indicating that it was generated by voltage-dependent Na^+^ channels. These results confirm that the properly chosen LC cells are functional and healthy neurons.

### Lack of Deleterious Effect of Fluo-3-AM Staining on Electrophysiological Responses

One consideration in carrying out the [Ca^2+^]c imaging experiments was that Ca^2+^ indicators, which bind Ca^2+^, could potentially affect the physiological [Ca^2+^]c dynamics, e.g. by completely chelating the free Ca^2+^ and thereby blocking the responses to caffeine. We tested whether Fluo-3-AM affects the physiology, using the caffeine-induced outward current as readout. Acutely dissociated LC neurons were treated with Fluo-3-AM, similarly as in [Ca^2+^]c imaging studies (Figures 11,12). However, the pretreatment did not modify the caffeine-induced outward currents (**Figure S2**). Thus, at least when used according to our protocol, Fluo-3 will detect [Ca^2+^]c changes without significantly perturbing Ca^2+^ dynamics.

### Decays of the [Ca^2+^]c Signals in Dendrites *vs.* Soma

In addition to the rise phase of the [Ca^2+^]c signals (Figure 11), another kinetic difference was noted among the subcellular regions. The fluorescence signal decayed in 2 out of 4 cells, and the decay was more rapid in the dendrites than in somatic regions (Figures S3A,B). The faster decay was clearly demonstrated by the traces whose peak amplitudes were normalized (Figure S3C, red arrowhead) and their slopes (Figure S3D, red arrowhead) (2 out of 4 cells). Thus both the rise (red arrows) and decay (red arrowheads) of the Ca^2+^ signal were faster in dendrites than in somata.

---

**Supplementary Figure 1.**
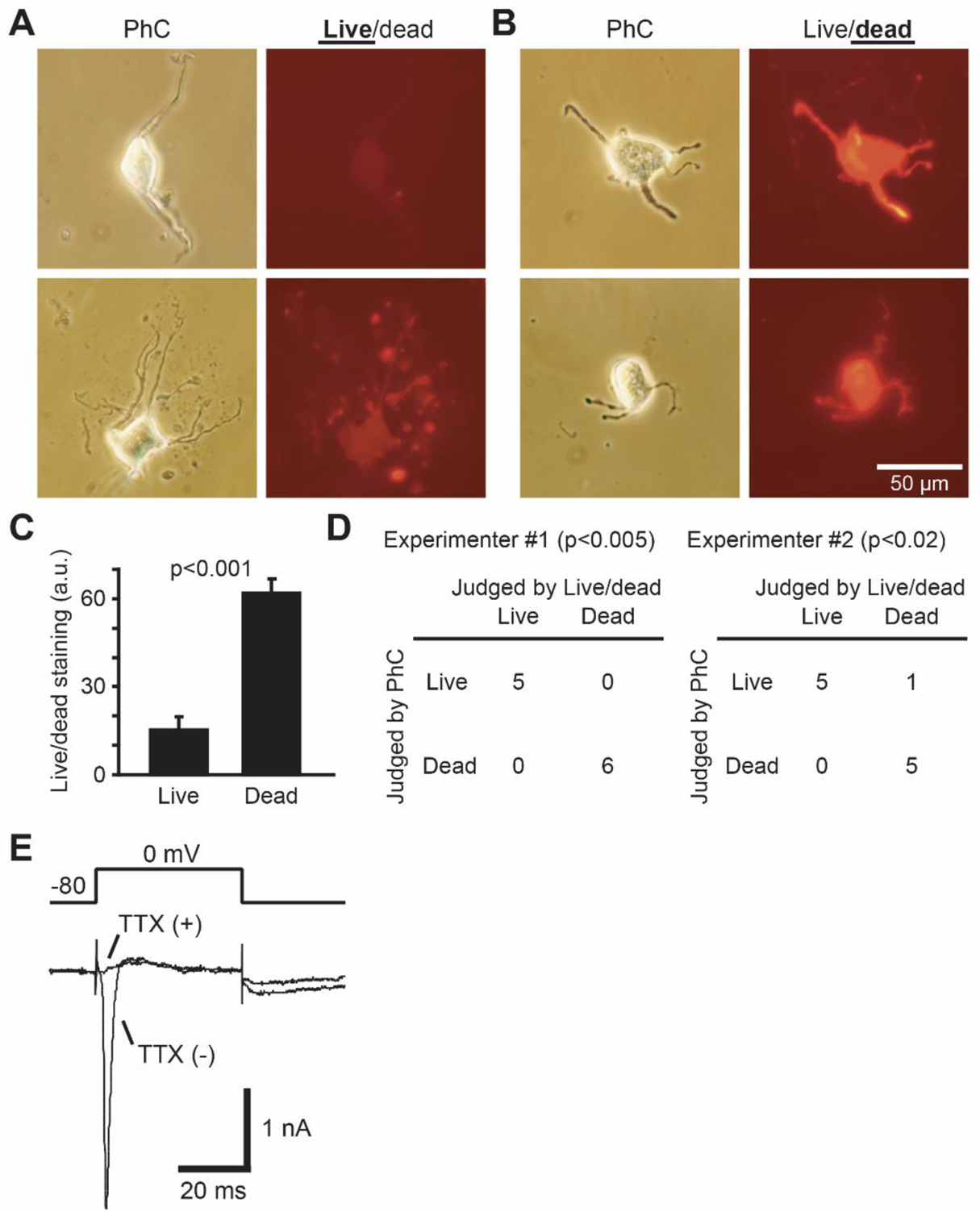
Live condition of the acutely dissociated locus coeruleus (LC) cells, as determined by fluorescent, live/dead staining **(A-D)** and electrophysiology **(E)**. **(A)** Phase-contrast (left, PhC) and fluorescent, live/dead staining (right) of two representative live cells, as evident from the low-intensity staining. **(B)** Phase-contrast (left) and fluorescent, live/dead staining (right) of two representative dead cells, as evident from the high-intensity staining. **(C)** Averaged fluorescence intensity of live/dead staining. The difference in staining intensity between cells determined to be live and dead was statistically significant (p<0.001, n=5 and 6 cells, t-test). Error bars represent the SEM. **(D)** Comparison of effectiveness of live/dead evaluation, by phase-contrast imaging (based on analysis by two experimenters, 1 and 2) *vs.* by live/dead staining. Numbers represent the number of cells categorized as live or dead. The match in the findings from the two approaches is statistically significant (p<0.005 and p<0.02 for two experimenters, respectively, Fisher’s exact probability test). **(E)** Dissociated LC cells were confirmed to be live and to be neurons based on the ability to induce of a voltage-dependent Na^+^ current, by holding the membrane potential at −80 mV and stepping to 0 mV for 40 msec. The current was blocked when 1 µM tetrodotoxin (TTX) was added to the solution.

**Supplementary Figure 2.**
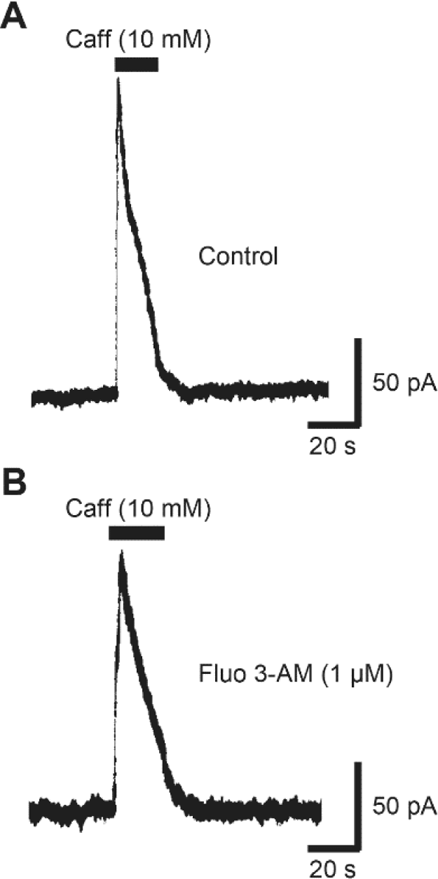
Lack of major effect of staining with Fluo-3-AM on electrophysiology of acutely dissociated LC neurons. The outward current induced by the application of 10 mM caffeine **(A)** was similar to that in a different cell pretreated with Fluo-3-AM **(B)** with the same protocol used for [Ca^2+^]c imaging. Holding potential was −44 mV as in the main figures.

**Supplementary Figure 3.**
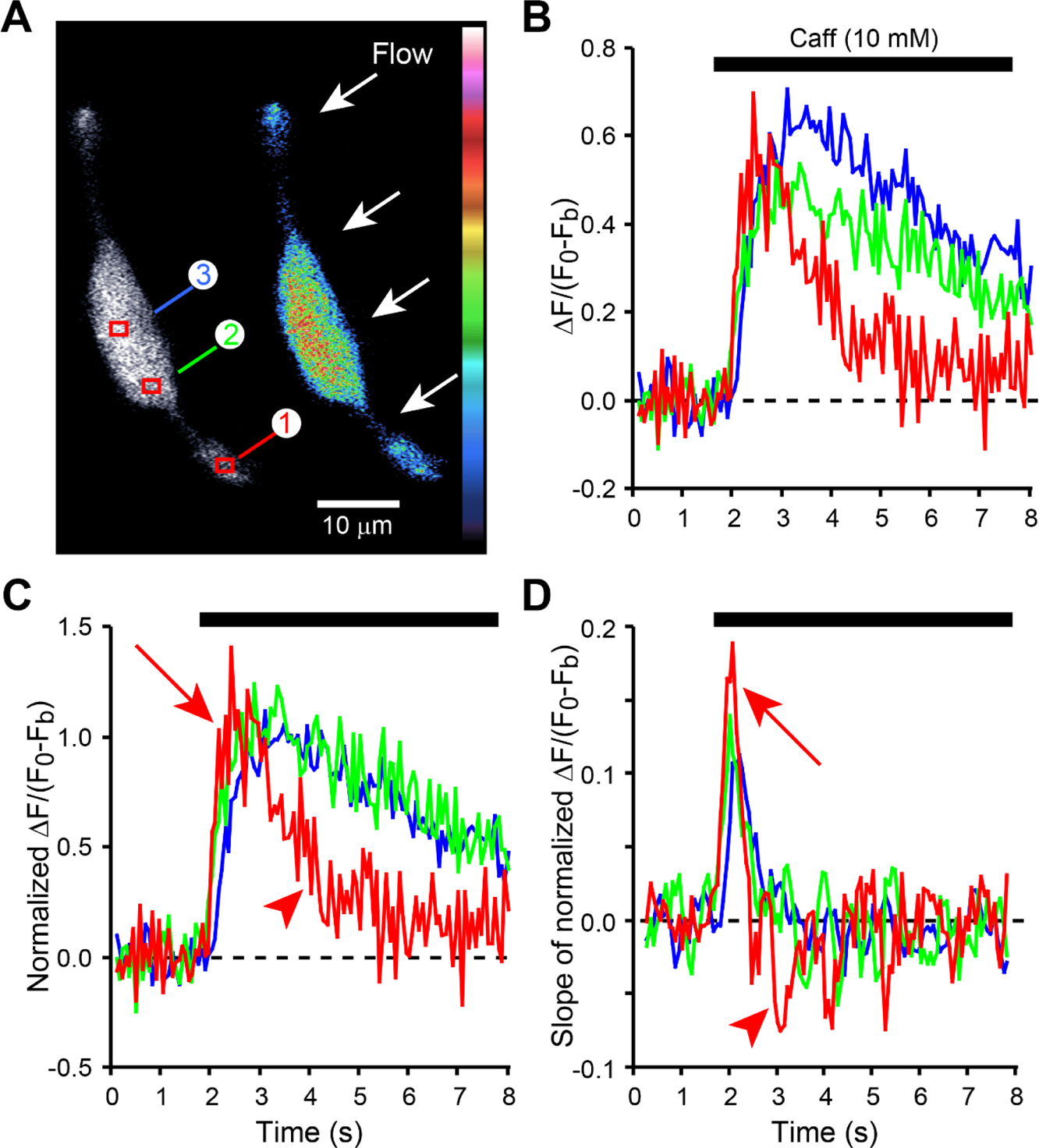
Decay of the caffeine-induced [Ca^2+^]c signal in a dendrite *vs.* a soma. Application of caffeine led to a [Ca^2+^]c increase (as in Figure 11), but in some neurons this was followed by a [Ca^2+^]c decrease, even in the continued presence of caffeine. **(A)** Location of each ROI. **(B)** All three ROIs, including one in a nuclear region (dark blue trace) showed a transient [Ca^2+^]c response, i.e. a rise followed by a decay. **(C)** In normalized traces, a dendrite showed more rapid rise (red arrow) and the decay (red arrowhead) than two regions in soma. **(D)** In a dendrite, the rates of both the rise (red arrow) and decay (red arrowhead) were higher than those in soma.

**Supplementary Figure 4.**
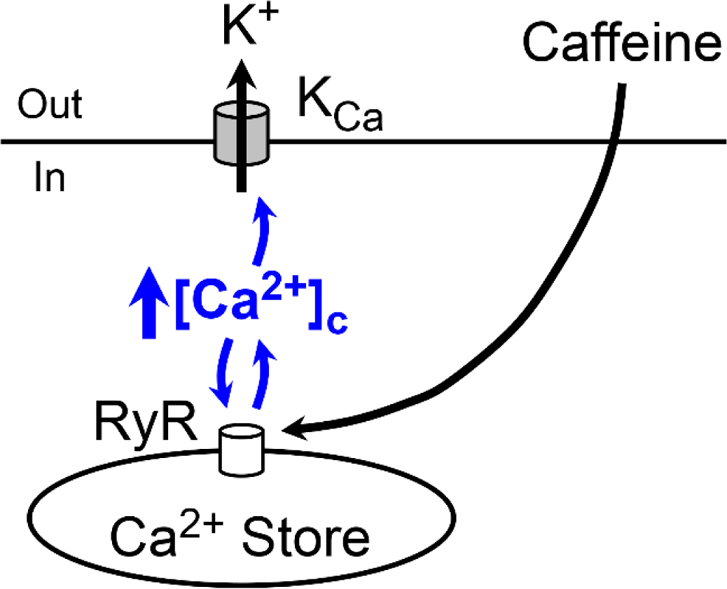
Summary diagram of the results reported in the current study, i.e. the impact of activating CICR in locus coeruleus neurons. **Abbreviations**: [Ca^2+^]c, cytosolic Ca^2+^ concentration; CICR, Ca^2+^-induced Ca^2+^ release; In, intracellular space; KCa, Ca^2+^-activated K^+^ channel of the non-small-conductance type (putatively, intermediate-conductance type); Out, extracellular space; RyR, ryanodine receptor.

